# Hyper-migratory CAR T cells alleviate ovarian cancer metastatic burden and improve prognosis

**DOI:** 10.1101/2025.09.28.679084

**Authors:** Praful R. Nair, Eban Hanna, Saurabh Joshi, Victoria Duarte-Alvarado, Challice L. Bonifant, Denis Wirtz

## Abstract

Cellular immunotherapy has shown remarkable efficacy in hematological malignancies but remains limited by infiltration issues in solid tumors, leading to poor treatment efficacy. We have recently shown that mesothelin-targeting CAR T cells co-transduced with cytokine-binding synthetic velocity receptors (VRs, referred to as CAR TV cells) demonstrate increased motility, improved infiltration in solid primary tumors, and lead to a better anti-tumor effect compared to CAR T cells that do not express VRs. However, it is metastasis that causes the vast majority of cancer related deaths and is difficult to target clinically, indicating an urgent unmet need. We show that these CAR T cells engineered to be hyper-migratory using VRs are highly effective against liver metastasis of ovarian cancer along different stages of the metastatic cascade. Mesothelin-targeting CAR TV cells expressing synthetic or native receptors responsive to the cytokine Interleukin-5 improved the survival of mice bearing an extremely high or ‘terminal’ level of metastatic burden compared to CAR T cells that did not express VRs. Against newly established metastatic lesions and lesions undergoing metastatic outgrowth, CAR TV cells showed a robust anti-cancer effect resulting in an improved prognosis compared to control CAR T cells. Histopathological assessments showed a substantial reduction of metastasis number and lesion size with CAR TV treatment, concomitant with increased immune cell infiltration in the metastatic regions. Our work demonstrates the efficacy of high-motility CAR T cells in a metastatic setting and extends their scope to the treatment of metastasis of solid tumors.

## Introduction

Despite the creation of several potent cellular immunotherapies, promising results observed in the laboratory have not always translated well to the clinic. This divide is particularly striking for solid tumors, where cellular immunotherapies have demonstrated underwhelming results (*1–3*). Most solid tumors feature a dense stromal collagenous matrix, that surrounds cancer cells, which, reduces the motility of immune cells trying to penetrate the tumor (*4*), in particular T cells (*5, 6*). One such desmoplastic solid tumor is ovarian cancer (*7, 8*), one of the most common gynecological malignancies and the fifth-leading cause of cancer mortality in women in the Unites States (*9*). The vast majority of cancer-related deaths are caused by metastasis, the spread of cancer cells from the primary tumor to secondary sites (*10, 11*). The majority of ovarian cancer patients are diagnosed with metastasis already present, and the five-year survival of patients with metastasis is ∼30% (*12*). This poor prognosis, combined with the limited improvement in therapies and frequent recurrence (*13, 14*), emphasizes that targeting metastatic ovarian cancer is the key to improving patient survival. Ovarian cancer spontaneously metastasizes to the liver and lungs, with the liver being the most common distant metastatic site (*15, 16*). Liver metastasis of ovarian cancer is particularly challenging to treat via immunotherapy due to its collagenous microenvironment (*17*) and a dearth of immune cells in the metastatic microenvironment (*18, 19*).

Chimeric antigen receptors (CARs) are engineered synthetic receptors that enable lymphocytes to recognize and attack cells expressing a specific target antigen (*20*). T cells expressing CARs (CAR T cells) that target cancer cells can elicit powerful anti-tumor responses. However, successes with CAR T treatment has been largely restricted to hematological malignancies, and its efficacy in solid tumors has been underwhelming (*20*). Solid tumors such as ovarian cancer, in which CAR T cells show poor infiltration (*21, 22*), comprises the vast majority of adult cancers (*9*) and represents a major obstacle as CAR T cells require direct physical contact with the cancer cells to invoke their anti-tumor effect (*20, 23*). Current designs of CAR T therapies have focused on enhanced specificity (*24–30*), reduced exhaustion (*31–35*) and enhanced activation (*36–39*), with little attention paid to improving the transport of these cells through the stromal matrix and their invasion into the tumor. The incorporation of this step can drastically decrease the observed immune cytotoxicity (*40*). The development of CARs that target mesothelin, a surface protein overexpressed on several solid tumors, has led to some successes against solid tumors (*41–43*), including ovarian cancer (*44, 45*). However, these mesothelin-targeting CAR T cells are still limited by inadequate infiltration in the tumor (*41, 42, 46–49*), which constitutes a major barrier to achieving a successful outcome.

We have previously shown that T cells secrete cytokines in a paracrine manner, that increases their motility in 3D collagen gels (*50*). Based on this observation, we designed velocity receptors (VRs) that are engineered to bind to the cytokines that drive sustained T cell migration, including interleukin-5 (IL-5) or interleukin-8 (IL-8). These VRs are either synthetic or simply an overexpression of the native cytokine receptor. Synthetic VRs comprises of four domains – binding, hinge, transmembrane, and signaling (*50*). The synthetic VRs use scFvs that bind to the cytokine of interest (IL-5 or IL-8) as the binding domain. The signaling domain combines receptor chains from IL5R (IL5RA and IL5RB) and TNFR (TNFR1, and TNFR2) in the configuration IL5RA-IL5RB-TNFR1-TNFR2. The hinge and transmembrane domain in the VRs correspond to that in CD8a. The expression of VRs on the surface of CAR T cells (referred to as CAR TV cells) increases cell motility by several-fold, enabling the CAR T cells to penetrate solid tumors much more effectively and in greater numbers. These cells retain the cytotoxicity of mesothelin CAR T cells, and combined with the increased tumor penetration, show a remarkable anti-tumor effect in primary tumors. However, despite the advances in treating primary tumors, targeting metastasis has been challenging in clinic (*51*), especially via cellular immunotherapy (*52*). Metastatic lesions also exhibit a desmoplastic collagenous stroma similar to that found in primary tumors (*17, 53, 54*). Hence, it stands to reason that the CAR TV cells, which we have shown to target and penetrate primary tumors *in vivo* would also provide benefit against metastatic burden. In particular, among the models we have tested previously (*50*), metastasis of ovarian cancer represents an enormous unmet need as described above.

Here, we show that highly motile mesothelin-targeting CAR (M5CAR) T cells expressing VRs are effective in reducing metastatic burden in mice bearing liver metastasis of ovarian cancer. CAR TV cells improved survival in three separate scenarios of metastatic burden: (i) extremely high metastatic burden corresponding to a couple of days of survival remaining, (ii) metastatic burden associated with actively proliferating metastatic colonies in the liver (metastatic outgrowth), and (iii) metastatic burden corresponding to newly established metastatic colonies in the liver (immediately post colonization). Three VRs were tested, and the overexpression of the native IL-5 receptor (referred to as V5-M5CAR) showed the best anti-metastatic potential, leading to a reduction in metastatic burden and/or survival benefit in every study across all three scenarios of metastasis. A synthetic VR comprised of an anti-IL-5 scFv and a signaling domain combining receptor chains from IL5RA, IL5RB, TNFR1, and TNFR2; (referred to as IL5-M5CAR) also showed promise, particularly in extending survival. These observations were supported by histopathological assessments of the liver, which showed fewer and smaller metastatic lesions with CAR TV treatment compared to CAR T. Importantly, these results were accompanied by greatly improved immune cell infiltration in the metastases, further supporting our hypothesis that the infiltration of immune cells forms a limiting barrier to CAR T cell therapy and that better immune cell infiltration would lead to a better treatment outcome.

Overall, our studies show that CAR T cells expressing velocity receptors show a robust anti-metastatic effect, reducing metastatic burden and improving survival across various levels of ovarian cancer metastatic burden in the liver. These results demonstrate the benefit of engineering hyper-migratory CAR T cells, that better penetrate metastatic lesions compared to other cellular immunotherapies.

## Results

### Establishment of liver metastasis of ovarian cancer

Ovarian cancer, one of the most common gynecological malignancies, spontaneously metastasizes to the liver and lungs among other sites (*16*). To create a metastasis model of ovarian cancer, we used an intravenous tail vein injection model, which we have previously used to establish metastasis models (*55*). Luciferase-P2A-mCherry OVCAR-3 cells were injected intravenously and their presence in the lungs was confirmed via bioluminescent IVIS imaging (Fig. S1 showing IVIS bioluminescence heatmap images). These mice were imaged daily to track the location and growth of metastatic lesions. Surprisingly, instead of colonizing the lung, the OVCAR-3 cells gradually migrated to the liver and established liver metastatic colonies. Immediately after the injection, OVCAR-3 cells reached the lungs and showed signal exclusively in the lungs till day 3, after which, luminescent signal was observed from both the lung and the liver (Fig. S1). This mixed signal was seen till day 6, with the liver signal growing stronger and the lung signal becoming weaker with time. From day 8 onwards, luminescent signal was only detected from the liver, indicating that OVCAR-3 cells complete liver colonization ∼7 days after injection.

The presence of liver metastases was confirmed upon the termination of the experiment, 26 days after the injection of OVCAR-3 cells. Excised livers showed extensive metastases, whereas the lungs appeared normal and did not exhibit any metastatic nodules. Examination of H&E stained sections showed numerous circular metastatic lesions with strong purple staining and irregular cellular organization (all indicative of cancerous growth, Fig. S2). Larger metastatic lesions showed lighter staining in the center, indicative of a necrotic core devoid of cells. To further confirm that these were indeed OVCAR-3 metastatic lesions, IHC was performed for the cytokeratins, CK8 and CK19, both of which are strongly expressed by OVCAR-3 cells (*56*) (Fig. S2). The areas identified as metastatic colonies in H&E sections showed intense brown staining for the two cytokeratins, confirming that these circular lesions in the liver were indeed OVCAR-3 metastatic colonies.

Hence, the tail-vein administration of OVCAR-3 ovarian cancer cells led to the formation of extensive metastatic colonies in the liver, which were verified by cytokeratin IHC.

### CAR TV cells improve the survival of mice bearing terminal metastatic burden

We have previously shown that mesothelin-targeting CAR (M5CAR) T cells that express velocity receptors (VRs) are highly motile in 3D environments (*50*) (Fig. 1a). These cells (abbreviated CAR TV cells) migrate three-times more compared to M5CAR T cells that did not express VRs, while retaining the cytotoxicity of the original M5CAR T cells. CAR TV cells showed dramatically increased infiltration into primary solid tumors, leading to potent anti-tumor effect *in vivo* (*50*). We hypothesized that this increased motility would also be effective against metastatic lesions that possess a similar collagenous desmoplastic matrix as primary tumors (*17, 53, 54*). Mesothelin is overexpressed in a wide range of solid cancers (*57*), is the subject of several ongoing CAR T cell clinical trials (*58*), and is an established target for CAR T cell therapy in ovarian cancer (*44, 45*). Since OVCAR-3 cells overexpress mesothelin and have been shown to be targeted by mesothelin-CAR T cells (*59*), mesothelin targeting CAR T (M5CAR T) cells were chosen as the baseline CAR T cell in this study.

**Figure 1.**
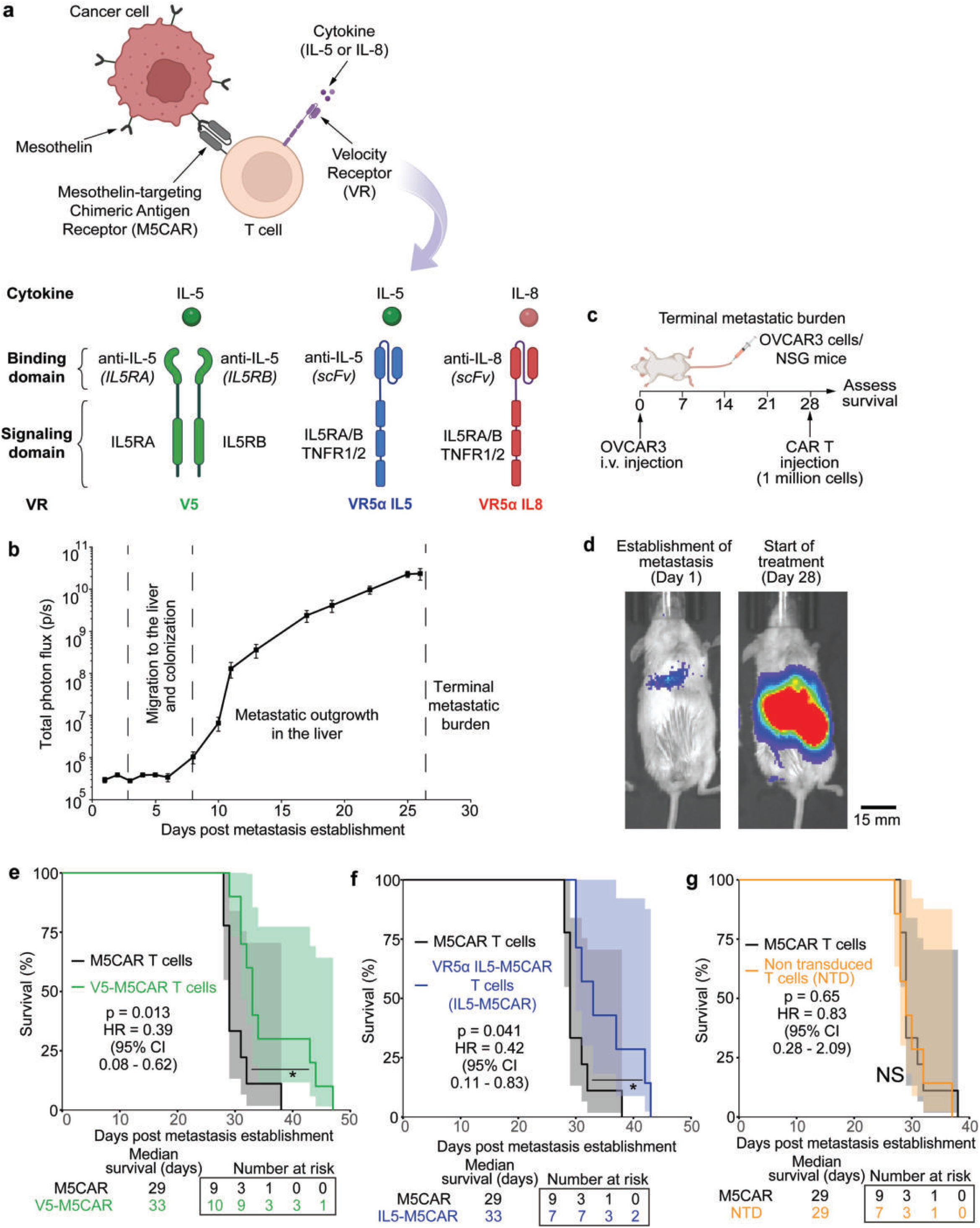
CAR TV therapy improves the survival of mice bearing terminal metastatic burden. **(a)** CAR TV cells express mesothelin-targeting chimeric antigen receptor (M5CAR) and velocity receptor (VR). M5CAR helps the T cells elicit an anti-tumor response when it binds to mesothelin, a surface protein which is overexpressed in several solid tumors. The VRs are responsive to cytokines that have been shown to drive T cell migration. The VRs used in this manuscript are responsive to the cytokines IL-5 or IL-8, and are either synthetic (VR5α IL5 and VR5α IL8) or an overexpression of the native IL-5 receptor (V5). Synthetic VRs comprise of an scFv targeting the cytokine of interest (IL-5 or IL-8) and a signaling domain combining receptor chains from IL5RA, IL5RB, TNFR, and TNFR2. **(b)** Growth of bioluminescent IVIS signal with time on a logarithmic scale shows the three relevant metastatic stages: colonization, metastatic outgrowth, and terminal metastatic burden. **(c)** NSG mice were injected with OVCAR-3 cells intravenously via a tail vein injection to form liver metastases. Four weeks after the injection, after a terminal metastatic burden was confirmed with IVIS signaling, mice were treated with 1 million M5CAR T cells expressing velocity receptors (VRs). M5CAR T cells that did not express VRs and non-transduced T (NTD) cells were the control. Survival was assessed following CAR T therapy. **(d)** IVIS luminescence signal after OVCAR-3 injection (day 1) and at the start of the treatment (day 28) showing extremely high metastatic burden. **(e)** V5-M5CAR T cell therapy significantly improves survival compared to M5CAR T cells not expressing VRs (median survival of 33 days versus 29 days, respectively, and hazard ratio of 0.39). Shaded area shows 95% confidence interval. **(f)** VR5α IL5-M5CAR T cell therapy significantly improves survival compared to M5CAR T cells (median survival of 33 days versus 29 days, respectively, and hazard ratio of 0.41). **(g)** Administration of non-transduced T cells does not lead to any survival benefit compared to M5CAR T cells. All data in this figure was generated with OVCAR-3 cells *in vivo* in NSG mice. p value: * p < 0.05, NS: Not Significant

Quantification of the temporal evolution of the metastatic burden shows two distinct phases of metastatic growth (Fig. 1b), corresponding to the final two steps of the metastatic cascade: colonization at the distant site and metastatic outgrowth (*60*). Metastatic lesions that were newly established in the liver and were yet to commence metastatic outgrowth were characterized by a low level of metastatic burden with a slow growth rate. As time elapses, the metastases that have progressed past colonization commence metastatic outgrowth, which is indicated by an intermediate level of metastatic burden with a high growth rate. The growth rate of metastatic burden (slope of the curve in Fig. 1b) is initially high during the metastatic outgrowth phase and eventually starts to plateau as the metastatic burden approaches an extensive level. This marks the transition to a third stage of metastatic burden, marked by an extremely high or ‘terminal’ level of metastasis.

The challenge of treating metastatic disease increases with higher metastatic burden (*61*). This third phase corresponding to late stage metastasis in terminal cancer patients is the most challenging to treat. To develop extensive metastatic burden in our model, the metastasis was allowed to grow for 28 days following OVCAR-3 injection (Fig. 1c). This time point results in extensive metastatic burden at the start of the treatment (visualized in Fig. 1d, ∼100,000 times the initial bioluminescent signal on day 1) and corresponds to ∼1-2 days prior to euthanasia as mandated by the ACUC protocol. These mice were treated with one of two CAR TV cells targeting the cytokine IL-5, V5-M5CAR T cells (which overexpress the native IL-5 receptor) and VR5α IL5-M5CAR T cells (which express an anti-IL-5 scFv and a signaling domain combining receptor chains from IL5RA, IL5RB, TNFR1, and TNFR2; referred to as IL5-M5CAR). The therapy dosage was fixed at 1 million cells per injection, which is much lower than the standard of 3-5 million cells/injection (*50*). Both CAR TV provided a significant survival benefit with a mean survival of 33 days compared to 29 days for M5CAR T cells which did not express VRs. This translated to a mean survival of five days post treatment for both CAR TV therapies versus a one day survival with M5CAR control (Fig. 1e-f). M5CAR T cells also did not provide an additional survival benefit compared to the administration of non-transduced T cells that did nor express CARs or VRs (NTD, Fig. 1g), potentially due to the lack of penetration of both the M5CAR T cells and NTD cells into the metastases, given that M5CAR T cells show vastly improved cancer cell killing *in vitro* compared to NTD cells (*62*).

Therefore, the administration of CAR TV cells expressing VRs provides a substantial survival benefit in mice bearing terminal metastatic burden compared to M5CAR T cell that do not express VRs.

### CAR TV cells reduce metastatic burden and provide survival benefit in mice bearing actively proliferating metastatic lesions

The success of CAR TV cells against terminal metastatic burden warrants an investigation into the efficacy of CAR TV cells against the earlier two stages of metastatic burden. To test the effectiveness of CAR TV cells against metastatic outgrowth in the liver, CAR TV treatment was administered two weeks post metastasis establishment (Fig. 2a). This timepoint corresponded to a metastatic burden that was visualized as less intense or widespread compared to terminal metastatic burden but substantially higher than that observed during initial establishment of metastasis (Fig. 2b, ∼1000 times the initial bioluminescent signal on day 1). The timepoint of 14 days post metastasis establishment was chosen as it lies near the beginning of the exponential growth phase (Fig. 1b), indicating that the metastatic colonies had finished colonization and had commenced outgrowth.

**Figure 2.**
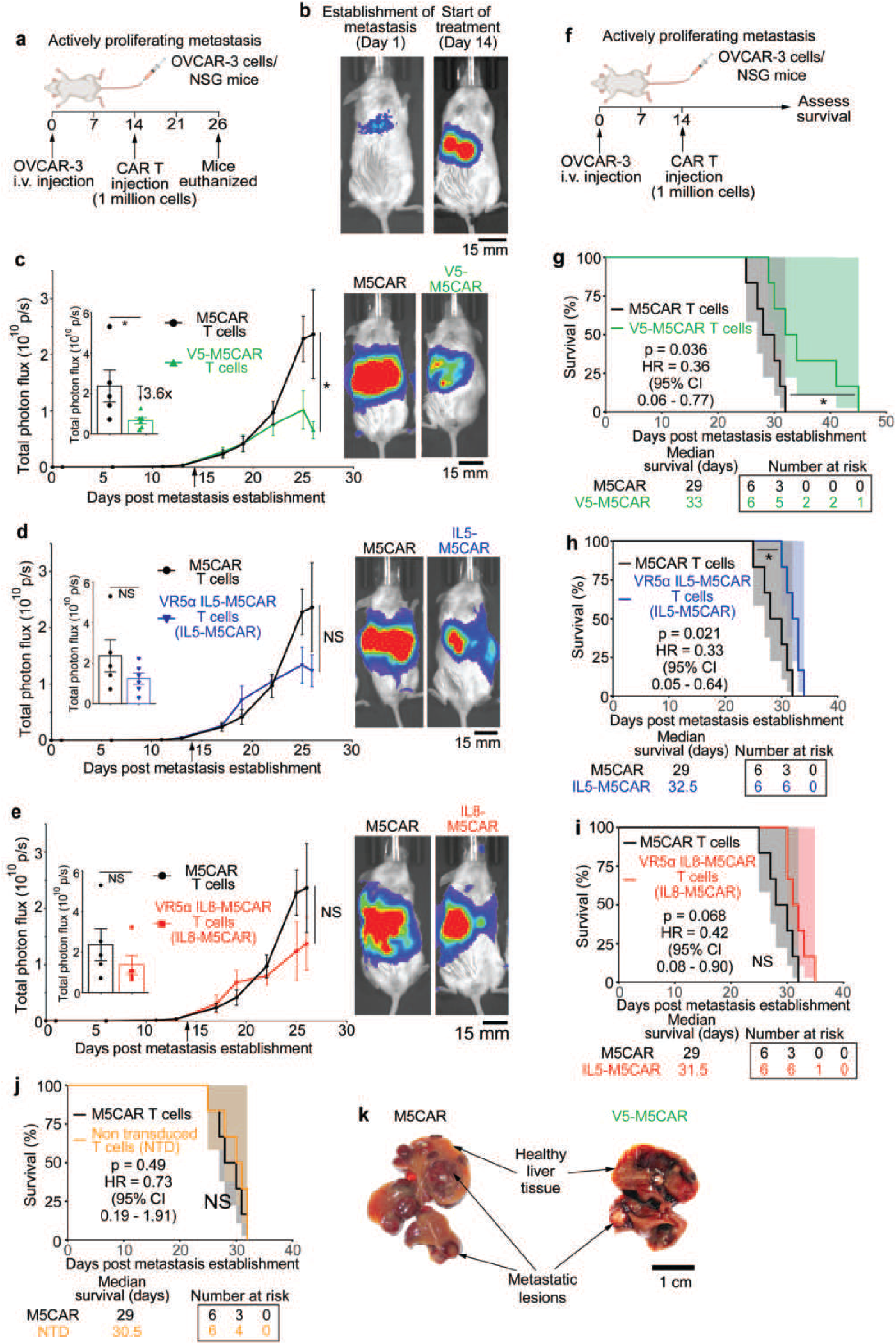
CAR TV therapy reduces metastatic burden and improves the survival of mice during metastatic outgrowth. **(a)** NSG mice were injected with OVCAR-3 cells intravenously via a tail vein injection to form liver metastases. Two weeks after injection, mice were treated with 1 million M5CAR T cells expressing velocity receptors (VRs). M5CAR T cells that did not express VRs were the control. Mice were euthanized 26 days after metastasis establishment. **(b)** IVIS luminescence signal after OVCAR-3 injection (day 1) and during the start of the treatment (day 14) showing an intermediate metastatic burden. **(c)** Administration of V5-M5CAR T cells led to a significant reduction in metastatic burden compared to M5CAR T cells that did not express VRs. **(d)** VR5α IL5-M5CAR and **(e)** VR5α IL8-M5CAR T cells reduced metastatic burden compared to M5CAR T cells, although they did not reach significance. **(f)** NSG mice were injected with OVCAR-3 cells intravenously via a tail vein injection to form liver metastases. Two weeks after injection, mice were treated with 1 million CAR T cells expressing velocity receptors (VRs). CAR T cells that did not express VRs and non-transduced T (NTD) cells were the control. Survival was assessed following CAR T therapy. **(g)** V5-M5CAR T cell therapy significantly improved survival compared to M5CAR T cells not expressing VRs (median survival of 33 days versus 29 days, respectively, and hazard ratio of 0.36). **(h)** VR5α IL5-M5CAR T cell therapy improves survival compared to M5CAR T cells (median survival of 32.5 days versus 29 days, respectively, and hazard ratio of 0.33). **(i)** Administration of VR5α IL8-M5CAR T cells or **(j)** non-transduced T cells do not lead to any survival benefit compared to M5CAR T cells. (k) Liver from CAR T cell treated mice show fewer and smaller metastatic lesions in the liver from mice treated with V5-M5CAR T cells compared to mice treated with M5CAR T cells that did not express VRs. All data in this figure was generated with OVCAR-3 cells *in vivo* in NSG mice. p value: * p < 0.05, NS: Not Significant

Administration of V5-M5CAR T cells decreased metastatic burden by ∼4-fold compared to mice treated with M5CAR T cells that did not express VRs (Fig. 2c, sample final mouse IVIS images on the right). IL5-M5CAR T cells also reduced metastatic burden compared to M5CAR T cells, however, it did not reach statistical significance (Fig. 2d). A similar result was observed with a third CAR TV cell type, VR5α IL8-M5CAR T cells, which express an anti-IL-8 scFv and a signaling domain combining receptor chains from IL5RA, IL5RB, TNFR1, and TNFR2 (referred to as IL8-M5CAR). As with IL5-M5CAR, IL8-M5CAR reduced metastatic burden, however, it was not statistically significant (Fig. 2e).

This pattern was also reflected in survival studies (Fig. 2f). Administration of V5-M5CAR and IL5-M5CAR significantly improved survival compared to M5CAR control (Fig. 2g-h, median survival of 33 days for V5-M5CAR, 32.5 days for IL5-M5CAR, and 29 days for V5-M5CAR). IL8-M5CAR improved the median survival (31.5 days vs. 29 days for M5CAR), however this improvement was not significant (Fig. 2i). As was the case with terminal metastatic burden, M5CAR T cells did not lead to an improvement in survival compared to non-transduced T cells that did not express CAR (Fig. 2j).

Hence, CAR TV administration provides therapeutic benefit against metastatic outgrowth. This benefit is evident in the reduced metastatic burden and the longer survival of mice receiving CAR TV treatment compared to M5CAR T cells.

### Anti-metastatic effect of CAR TV cells is evident in the histopathology of livers

To better understand the impact of CAR TV cells on liver metastases of ovarian cancer, the livers of all mice from the fixed timeline study (Fig. 2c-e) were excised. Mice treated with V5-M5CAR T cells showed fewer metastatic lesions that were smaller in size compared to mice receiving M5CAR T cell therapy (Fig. 2k, full panel of livers in Fig. S4b). H&E stains of liver sections were examined to better characterize the histological impact of CAR TV treatment on liver metastasis. H&E liver sections from mice treated with M5CAR T cells showed multiple large metastatic lesions per section, with several being large enough to possess necrotic cores (Fig. 3b, full panel of livers in Fig. S5). V5-M5CAR T cell treated mice, in contrast, had smaller and fewer metastatic lesions, with very few being large enough to have necrotic cores, consistent with the alleviation of metastatic burden in Fig. 2c. Other CAR TV therapies (IL5-M5CAR and IL8-M5CAR) showed histological characteristics on the spectrum between V5-M5CAR T cells and M5CAR T cells that did not express VRs.

**Figure 3.**
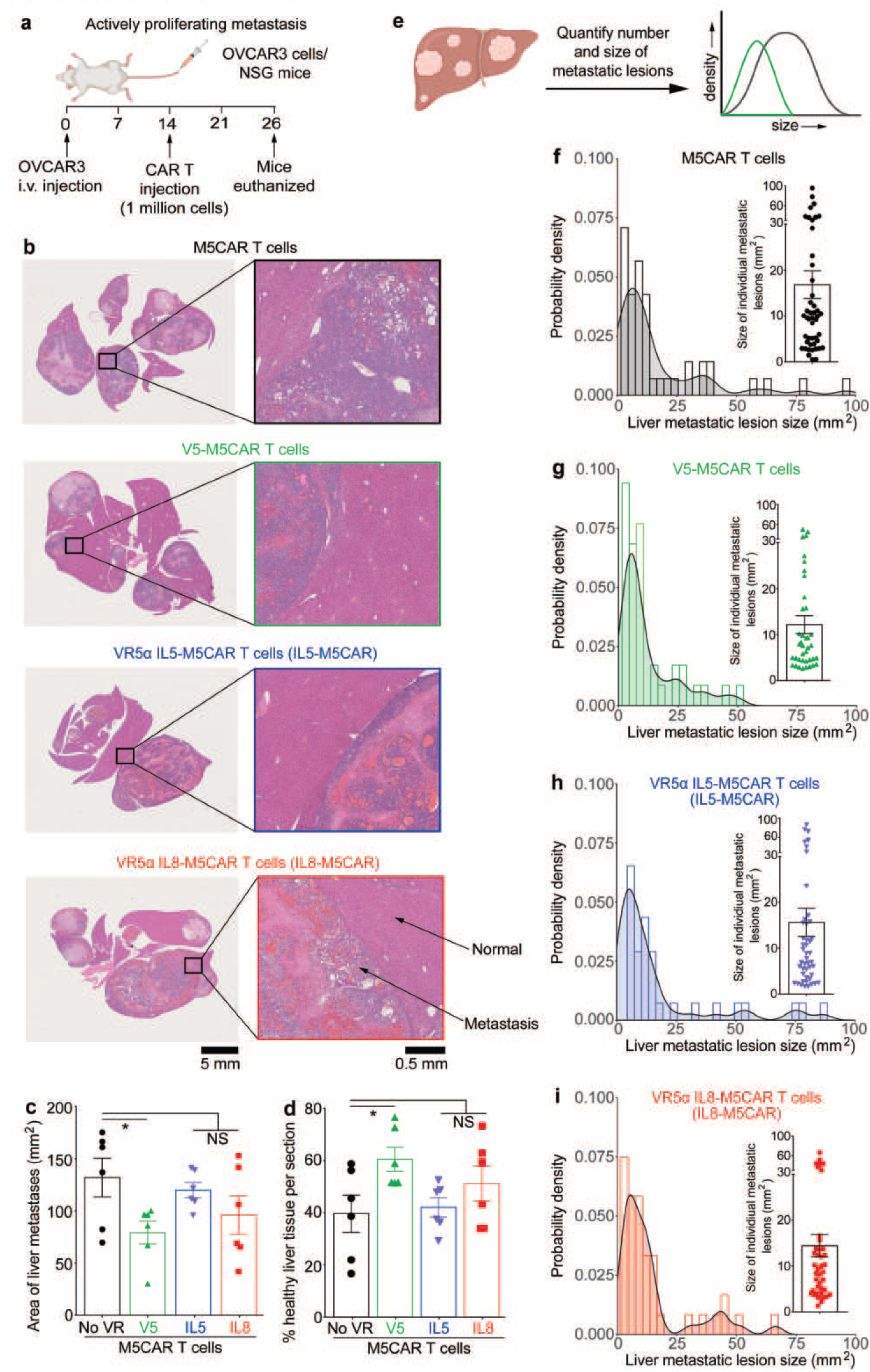
CAR TV treatment reduces liver metastatic burden of ovarian cancer when administered against metastatic outgrowth. **(a)** NSG mice were injected with OVCAR-3 cells intravenously via a tail vein injection to form liver metastases. Two weeks after injection, mice were treated with 1 million M5CAR T cells expressing velocity receptors (VRs). M5CAR T cells that did not express VRs were the control. Mice were euthanized 26 days after metastasis establishment. **(b)** H&E staining of liver sections show that mice treated with M5CAR T cells showed numerous large metastatic circular lesions characterized by intense purple staining, irregular cell arrangement, and a necrotic core in the case of larger lesions (indicated by lighter staining in the middle). Administration of V5-M5CAR T cells reduced the size and number of these metastatic lesions, indicating an overall decrease in metastatic burden. **(c)** Quantification of total metastatic burden per liver showed that V5-M5CAR T cell treated mice had the lowest liver metastatic burden. **(d)** Quantification of percentage of healthy liver tissue (defined as areas exhibiting normal liver cellular architecture) showed that a CAR TV treated mice exhibited livers with a greater healthy tissue fraction. **(e)** Histogram of metastatic lesion sizes were examined to better understand the impact of CAR TV cell therapy on individual metastatic lesions. **(f-g)** V5-M5CAR T cell therapy not only reduced the number of metastases, but were particularly effective in eliminating larger metastatic lesions, evident in the lack of a long right-hand side tail in the V5-M5CAR histogram (green) compared to a long-tailed distribution for M5CAR control (black). **(h-i)** Administration of IL5-M5CAR and IL8-M5CAR did not appreciably alter the histogram of metastatic lesion size but showed a shortening of the histogram tail compared to M5CAR T cell treated livers. All data in this figure was generated with OVCAR-3 cells *in vivo* in NSG mice. p value: * p < 0.05, NS: Not Significant

To connect these histological features with the observed decrease in metastatic burden and longer survival in Fig. 2, the metastatic burden, healthy tissue area, and the size distribution of metastases were calculated. Quantification of total area of metastatic lesions per slide across all mice showed that CAR TV treatment led to a decrease in total metastatic burden, with V5-M5CAR treatment being the most effective (Fig. 3c). This was accompanied by a corresponding increase in healthy liver tissue exhibiting structured cellular organization (Fig. 3d). Greater healthy liver tissue fraction, along with the reduced metastatic burden, would result in less impairment of liver function, which could partially explain the lack of signs of distress and longer survival in mice receiving CAR TV treatment, particularly for V5-M5CAR therapy. Histograms of metastasis sizes show striking differences between the size distributions of metastatic colonies in mice receiving CAR TV therapy versus M5CAR cells that do not express VRs. M5CAR treated livers showed a positively-skewed normal distribution with a long tail, indicating the presence of several large metastatic lesions (Fig. 3f). V5-M5CAR treatment dramatically shortened this tail, indicating that V5-M5CAR therapy was effective in preventing the incidence of large metastatic foci, especially metastatic foci larger than 50 mm^2^ (Fig. 3g). Consistent with previous results in Fig. 2, IL5-M5CAR (Fig. 3h) and IL8-M5CAR (Fig. 3i) led to size distributions with longer tails that resembled M5CAR distribution more closely than V5-M5CAR distributions. Finally, since the initial destination of the injected OVCAR-3 cells were the lungs, H&E lung sections were examined for metastatic colonies. Lung H&E sections showed no major metastatic colonies (Fig. S6), consistent with the permanent disappearance of bioluminescent signal from the lungs roughly a week after OVCAR-3 injection.

In sum, V5-M5CAR T cell therapy led to the lowest metastatic burden and the greatest survival benefit. This result was supported by liver sections, which showed smaller and fewer metastatic lesions with V5-M5CAR treatment as well as greater healthy liver tissue fraction.

### CAR TV cells show extensive infiltration in metastatic lesions

In order to verify that the increased efficacy of CAR TV cells was indeed due to better metastasis infiltration, we quantified the number of individual immune cells in the metastatic lesions using digital pathology. Digital pathology was preferred for its speed, unbiased nature, and reproducibility compared to manual detection via a pathologist (*63–65*). CellViT++ (*66, 67*), a transformer-based model trained on liver samples, was leveraged for cell segmentation and classification into cell types (Fig. 4a). V5-M5CAR treated metastases showed extensive immune cell infiltration, while M5CAR treated lesions only exhibited a few immune cells (Fig. 4b, immune cells outlined in green). Quantification of this immune infiltration showed that CAR TV treatment led to a ∼4-fold increase in immune cell infiltration in the metastatic lesions compared to M5CAR T cells (Fig. 4c). V5-M5CAR T cells, which had the greatest impact on metastatic burden and the best improvement in survival, showed the highest infiltration, followed by IL5-M5CAR and IL8-M5CAR. To better understand the impact of immune cell infiltration on the anti-metastatic activity of CAR T cells, we plotted the metastatic burden on the final day against immune cell infiltration (Fig. 4d). The final metastatic burden showed an inverse linear correlation with immune cell infiltration with a high goodness-of-fit (R^2^ = 0.96). Across all CAR T treatments, better infiltration correlated with lower final metastatic burden. Similar trends were seen with survival following CAR TV treatment (Fig. 4e-f). Increased infiltration of CAR TV cells was also correlated with better survival post treatment (Fig. 4e, R^2^ = 0.92) and higher median survival (Fig. 4f, R^2^ = 0.99).

**Figure 4.**
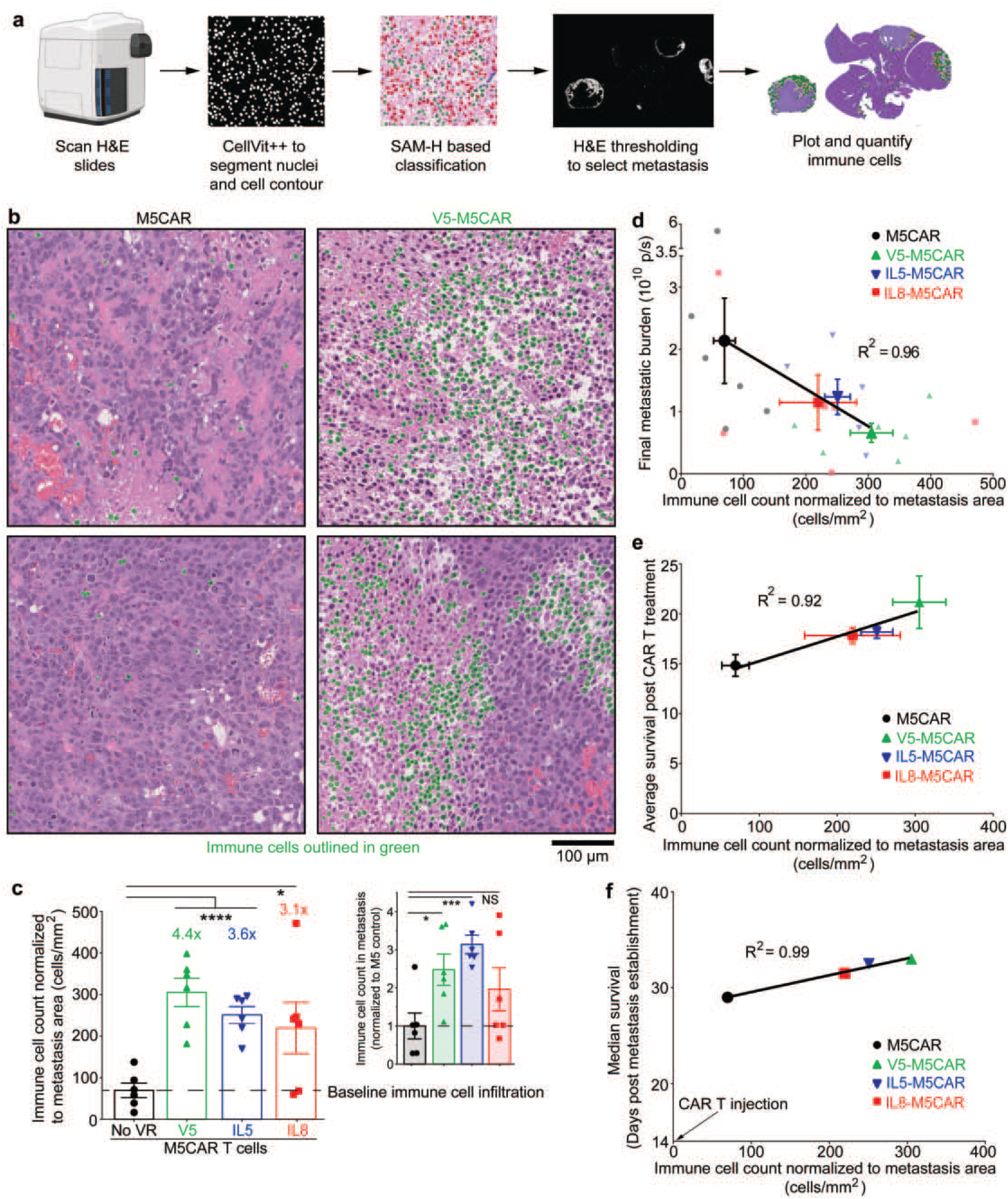
CAR TV cells show extensive infiltration in liver metastasis of ovarian cancer. **(a)** Workflow to quantify immune cell infiltration in liver metastasis from H&E slides using CellViT++. **(b)** Representative images showing immune cell infiltration (green) in liver metastasis for M5CAR and V5-M5CAR treated livers. M5CAR treated livers show only a few immune cells (outlined in green), while liver metastases treated with V5-M5CAR show extensive immune cell infiltration inside the metastatic lesions. **(c)** CAR TV cells showed increased infiltration compared to M5CAR T cells that did not express VRs. The highest level of immune cell infiltration (normalized to metastasis area) was observed with V5-M5CAR T cells, which also showed the greatest anti-metastatic efficacy and the best improvement in survival (Fig. 2). Immune infiltration quantification not adjusted for metastasis area is shown in the inset. CAR TV treated livers still show increased immune cell infiltration, despite having a smaller metastasis area. **(d)** Inverse correlation between immune cell infiltration and final metastatic burden. Increased immune cell infiltration with CAR TV administration was associated with decreased IVIS luminescent signal on the final day (linear fit, R^2^ = 0.96). **(e)** Increased infiltration of CAR TV cells was also correlated with better survival, as seen with average survival post treatment (linear fit, R^2^ = 0.92), and **(f)** median survival (linear fit, R^2^ = 0.99). All data in this figure was generated by quantification of OVCAR-3 metastasis *in vivo* in NSG mice. p value: * p < 0.05, *** p < 0.001, **** p < 0.0001, NS: Not Significant

Hence, increased CAR T cell infiltration was correlated with better anti-metastatic performance *in vivo*.

### CAR TV therapy is effective in mice bearing newly established metastatic colonies

Given the efficacy of CAR TV cells against higher metastatic burden, we sought to assess the impact of CAR TV cells on newly established metastatic colonies that had not yet started metastatic outgrowth. This model was established by administering the CAR TV injection one week after the initial OVCAR-3 injection (Fig. 5a). This timepoint was chosen as it gives enough time for the signal to shift to the liver and form nascent metastatic colonies (Fig. S1), is still early enough to represent a low metastatic burden (Fig. 5b, ∼10 times the initial bioluminescent signal on day 1), and it corresponds to a low growth rate of metastatic burden, evidenced by a nearly flat growth rate (Fig. 1b).

**Figure 5.**
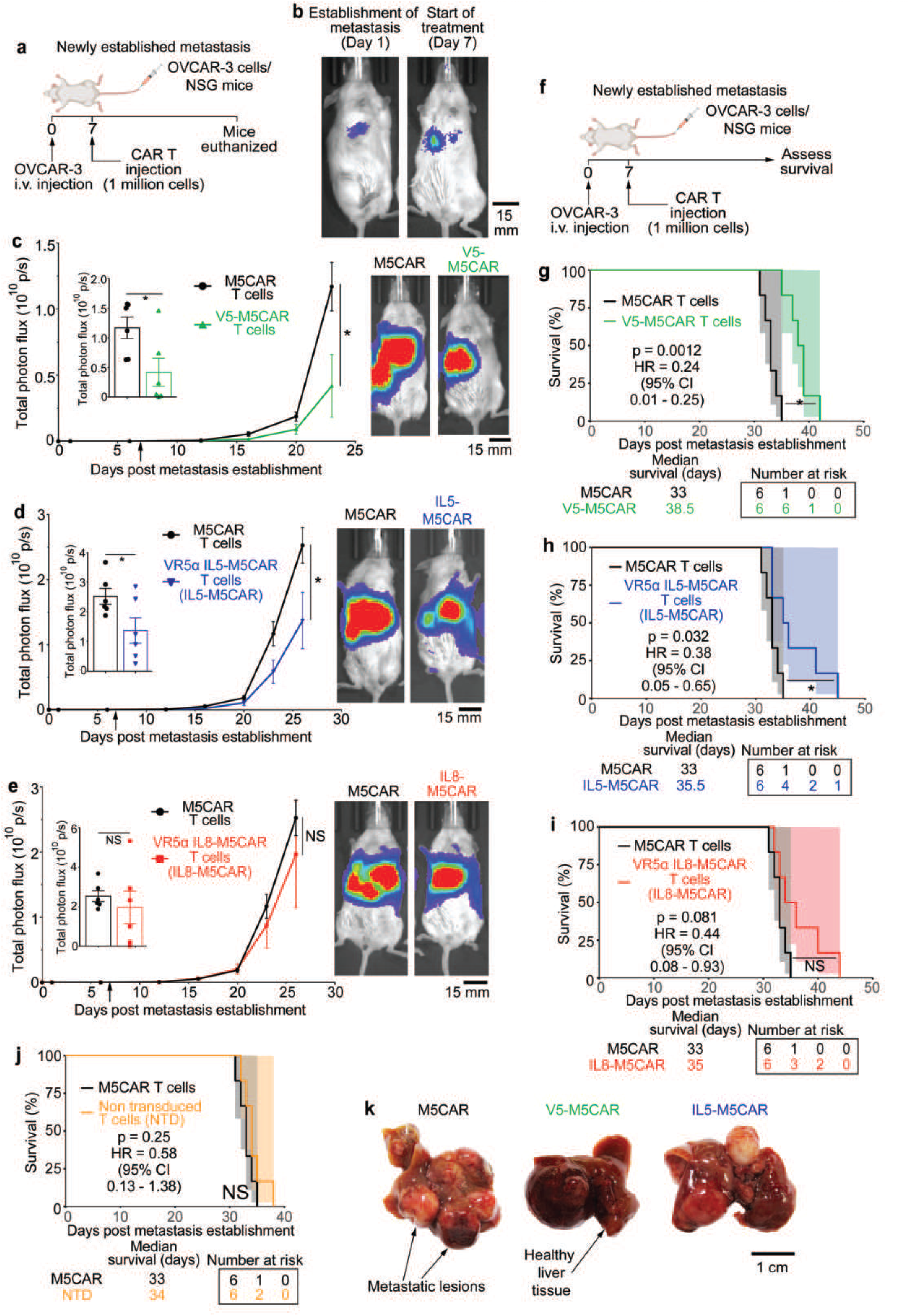
CAR TV therapy reduces metastatic burden and improves the survival of mice bearing newly established metastatic lesions. **(a)** NSG mice were injected with OVCAR-3 cells intravenously via tail vein injection to form liver metastases. One week after the injection, mice were treated with 1 million M5CAR T cells expressing velocity receptors (VRs). M5CAR T cells that did not express VRs were the control. Mice were euthanized at a predetermined time point. **(b)** IVIS luminescence signal after OVCAR-3 injection (day 1) and during the start of the treatment (day 7) showing a low metastatic burden. **(c)** Administration of V5-M5CAR T cells led to a nearly three-fold decrease in metastatic burden compared to M5CAR T cells that did not express VRs. **(d)** VR5α IL5-M5CAR administration led to a significant decrease in metastatic burden compared to M5CAR T cells. **(e)** VR5α IL8-M5CAR T cells reduced metastatic burden compared to M5CAR T cells, although they did not reach significance. **(f)** NSG mice were injected with OVCAR-3 cells intravenously via a tail vein injection to form liver metastases. One week after injection, mice were treated with 1 million CAR T cells expressing velocity receptors (VRs). CAR T cells that did not express VRs and non-transduced T (NTD) cells were the control. Survival was assessed following CAR T therapy. **(g)** V5-M5CAR T cell therapy significantly improved survival compared to M5CAR T cells not expressing VRs (median survival of 38.5 days versus 33 days, respectively, and hazard ratio of 0.24). **(h)** VR5α IL5-M5CAR T cell therapy significantly improves survival compared to M5CAR T cells (median survival of 35.5 days versus 33 days, respectively, and hazard ratio of 0.38). **(i)** Administration of VR5α IL8-M5CAR T cells or **(j)** non-transduced T cells do not lead to any survival benefit compared to M5CAR T cells. **(k)** Livers from V5-M5CAR and IL5-M5CAR T cell treated mice show fewer and smaller metastatic lesions in the liver from mice treated with V5-M5CAR T cells compared to mice treated with M5CAR T cells that did not express VRs. All data in this figure was generated with OVCAR-3 cells *in vivo* in NSG mice. p value: * p < 0.05, NS: Not Significant

As had been observed previously, V5-M5CAR T cells performed the best, leading to a three-fold decrease in metastatic burden compared to M5CAR T cells that did not express VRs (Fig. 5c). IL5-M5CAR T cells, which led to survival benefit with terminal metastasis and metastatic outgrowth, also significantly reduced final metastatic burden (Fig. 5d). IL8-M5CAR T cells also showed a modest reduction in metastatic burden compared to M5CAR T cells, however, this result did not reach statistical significance (Fig. 5e). These trends were also reflected in survival studies (Fig. 5f). Both V5-M5CAR (median survival 38.5 days, Fig. 5g) and IL5-M5CAR (median survival 35.5 days, Fig. 5h) led to an improvement in survival and a corresponding decrease in hazard ratios compared to M5CAR T cells that did not express VRs (median survival 33 days). IL8-M5CAR T cells (median survival 35 days) and NTD cells (median survival 34 days) did not significantly change survival compared to M5CAR T cells. (Fig. 5i-j)

Hence, CAR TV cells are effective in reducing metastatic burden across different stages of the metastatic cascade, corresponding to multiple levels of metastatic burden. Administration of CAR TV cells, particularly V5-M5CAR and IL5-M5CAR T cells that are responsive to the cytokine IL-5, led to a substantial reduction in metastatic burden and improved survival.

### Liver sections validate the anti-metastatic effect of CAR TV cells on newly-established metastatic lesions

In order to study the impact of CAR TV treatment on newly established metastases, the livers were excised and subjected to further histopathological analysis as described previously. CAR TV treated livers were morphologically more normal than M5CAR livers, showing more healthy tissue and far fewer metastatic lesions (Fig. 5k, full panel of livers in Fig. S7b). V5-M5CAR treated livers were the healthiest with few metastatic lesions across the full panel of livers (Fig. S7b). Liver H&E sections showed extensive metastatic colonization for mice receiving M5CAR or IL8-M5CAR treatments, but not in mice receiving V5-M5CAR or IL5-M5CAR treatment (Fig. 6b, full panel of livers in Fig. S8). Quantification of liver metastatic burden showed that V5-M5CAR therapy led to the lowest metastatic burden, with IL5-M5CAR treatment also significantly alleviating metastatic burden (Fig. 6c). This was accompanied by the highest healthy liver tissue fraction for these two CAR TV treatments (Fig. 6d). Histogram of metastatic lesion size showed that V5-M5CAR treatment was effective at preventing the occurrence of larger metastatic lesions, which were prevalent in livers treated with M5CAR T cells that did not express VRs (Fig. 6e-f). IL5-M5CAR led to a slight decrease in the right-hand tail of the distribution along with a decrease in the total number of metastases (Fig. 6g), while IL8-M5CAR livers showed a similar size range compared to M5CAR treated livers (Fig. 6h). Once again, lung H&E sections showed no metastatic foci (Fig. S9).

**Figure 6.**
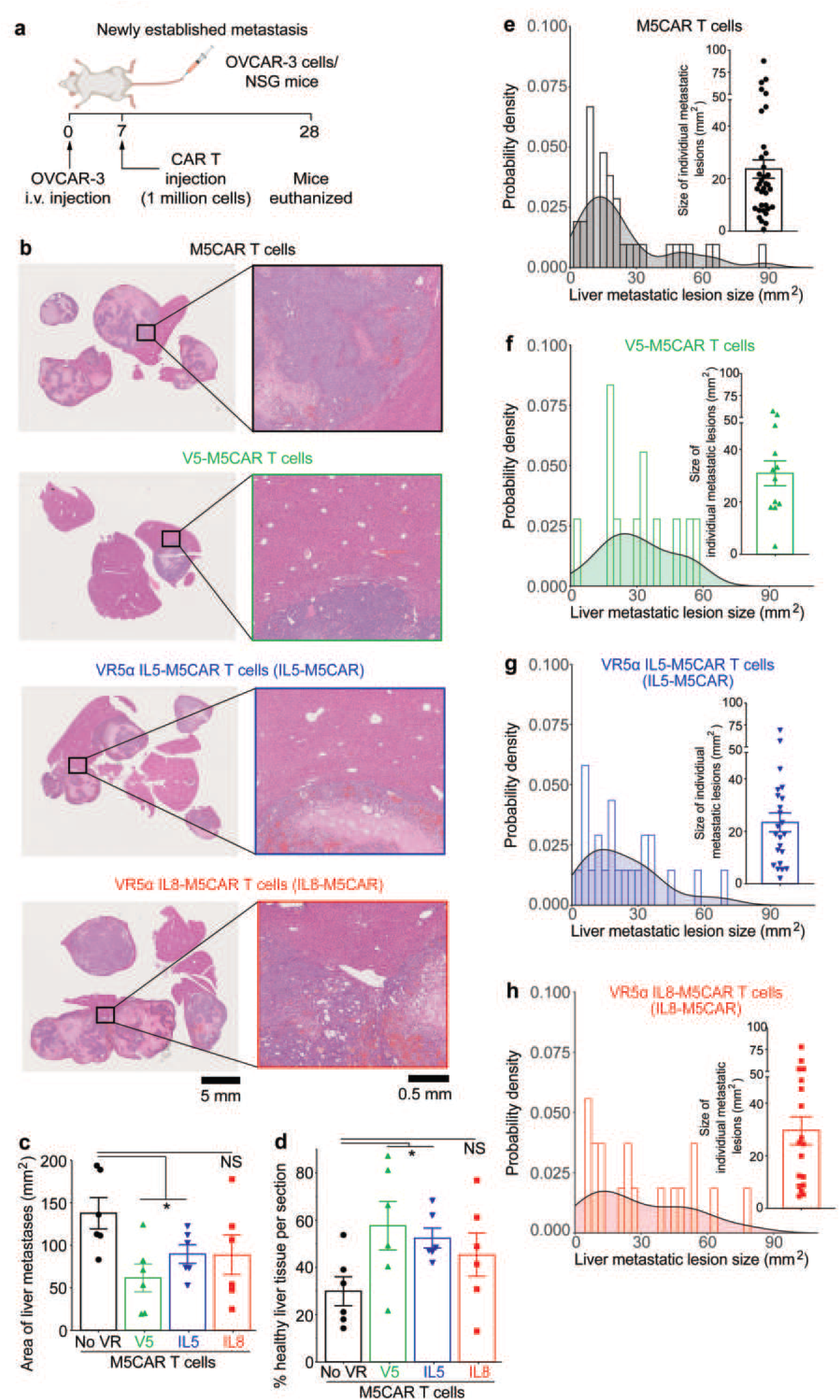
CAR TV treatment reduces liver metastatic burden of ovarian cancer when administered immediately post metastatic colonization. **(a)** NSG mice were injected with OVCAR-3 cells intravenously via tail vein injection to form liver metastases. One week after the injection, mice were treated with 1 million M5CAR T cells expressing velocity receptors (VRs). M5CAR T cells that did not express VRs were the control. Mice were euthanized at a predetermined time point. **(b)** H&E staining of liver sections show that mice treated with V5-M5CAR or IL5-M5CAR T cells showed a reduction in the size and number of metastatic lesions, consistent with the decrease in metastatic burden observed via IVIS imaging. **(c)** Quantification of total metastatic burden per liver showed that V5-M5CAR and IL5-M5CAR T cell treated mice exhibited significantly reduced liver metastatic burden. **(d)** Quantification of percentage of healthy liver tissue showed an increase in the healthy tissue fraction for CAR TV treated mice. **(e-h)** CAR TV cell therapy was effective in preventing large metastatic foci, with V5-M5CAR (green) and IL5-M5CAR (blue) being particularly effective in eliminating larger metastatic lesions, compared to M5CAR control (black, long-tailed distribution). Administration of IL8-M5CAR did not appreciably alter the histogram of metastatic lesion size but showed a shortening of the histogram tail compared to M5CAR control. All data in this figure was generated with OVCAR-3 cells *in vivo* in NSG mice. p value: * p < 0.05, NS: Not Significant

In sum, our hyper-migratory CAR T cells are effective against liver metastasis of ovarian cancer. CAR TV cells reduced metastatic burden and improved the survival of mice bearing various levels of metastatic burden. CAR T cells overexpressing the native IL-5 receptor (referred to as V5-M5CAR) were the most effective, followed by CAR T cells which expressed an anti-IL-5 scFv and a signaling domain combining receptor chains from IL5RA, IL5RB, TNFR1, and TNFR2 (referred to as IL5-M5CAR). Concomitant with the reduction in metastatic burden, CAR TV treatment led to a profound improvement in liver histology and was associated with higher immune cell infiltration in the metastatic lesions. Our results extend the scope of CAR TV cells from primary tumors (*50*) to cancer metastasis, the primary cause of cancer-related deaths.

## Discussion

Here, we demonstrate the advantages of engineered hyper-migratory CAR T cells over CAR T cells that do not express velocity receptors (VRs). CAR T cells expressing VRs led to lower metastatic, burden, longer survival, histopathologically healthier livers, and better infiltration in the metastases than CAR T cells that do not express VRs. CAR T cells have suffered from poor penetration in solid tumors (*21, 22*) and this study – along with our other efforts - demonstrates the benefit of using CAR TV cells over CAR T cells. One of the strengths of our approach is its orthogonality, which leads to full compatibility with other CAR constructs. We have demonstrated a proof-of-concept study using a mesothelin-targeting CAR. However, VRs can be used in parallel with other efforts to enhance specificity and reduce fast exhaustion of CAR T therapy (*22, 24–39*) to complement these studies.

Another notable detail is the low dosage of cells used in treatments. Low levels of immune cell infiltration in solid tumors with cellular immunotherapy has necessitated higher cellular dosage in those these treatments (*1–3, 50*). This could exacerbate the side-effects associated with CAR T therapy such as cytokine release syndrome, immune effector cell-associated neurotoxicity syndrome, and cytopenias (*68*). Given the higher infiltrative capacity of CAR TV cells, a lower dosage of CAR T cell was chosen due to the potential to reduce adverse off-target effects in future studies and clinical trials, and make these adverse events easier to manage. All CAR T treatments in this manuscript involved an injection of 1 million cells, which is considerably lower than the standard 3-5 million cells per injection in solid tumors (*50, 69–72*). Reduced number of cells would also be easier, faster, and potentially cheaper to produce. This low dosage could also explain the lack of significance seen with IL8-M5CAR T cells despite seeing a decrease in metastatic burden and extension of survival, accompanied by increased infiltration of IL8-M5CAR T cells compared to M5CAR T cells without VRs. A standard 3 million cell injection or a longer experimental timeframe could have led to a significant reduction in metastatic burden with IL8-M5CAR T cells.

Our results also show that not all VRs were equally effective. V5-M5CAR (which overexpresses the native IL-5 receptor) led to the best results and showed the greatest infiltration in the metastatic lesions. IL5-M5CAR (which expresses an anti-IL-5 scFv and a signaling domain combining receptor chains from IL5RA, IL5RB, TNFR1, and TNFR2) also showed robust immune cell infiltration, which translated to an anti-metastatic performance comparable to V5-M5CAR. IL8-M5CAR (which expressed an anti-IL-8 scFv and a signaling domain combining receptor chains from IL5RA, IL5RB, TNFR1, and TNFR2) does show penetration but does not work well, which could be due to a number of factors as discussed in (*50*). While penetration is a critical part of the CAR T cell-based anti-tumor response, an in-depth understanding of the impact of VR expression on other aspects of CAR T cell function including cytotoxicity, activation, and exhaustion in future studies will help better understand the differences observed between the different CAR TVs. Finally, this study utilized the three most promising VR constructs from our previous work (*50*) based on their *in vivo* performance in primary tumors, including ovarian cancer. Given the modular architecture of velocity receptor, there are well over 30,000 possible VR combinations and our future work will evaluate more combinations to identify better VRs.

## Supplementary Figure

**Supplementary Figure 1.**
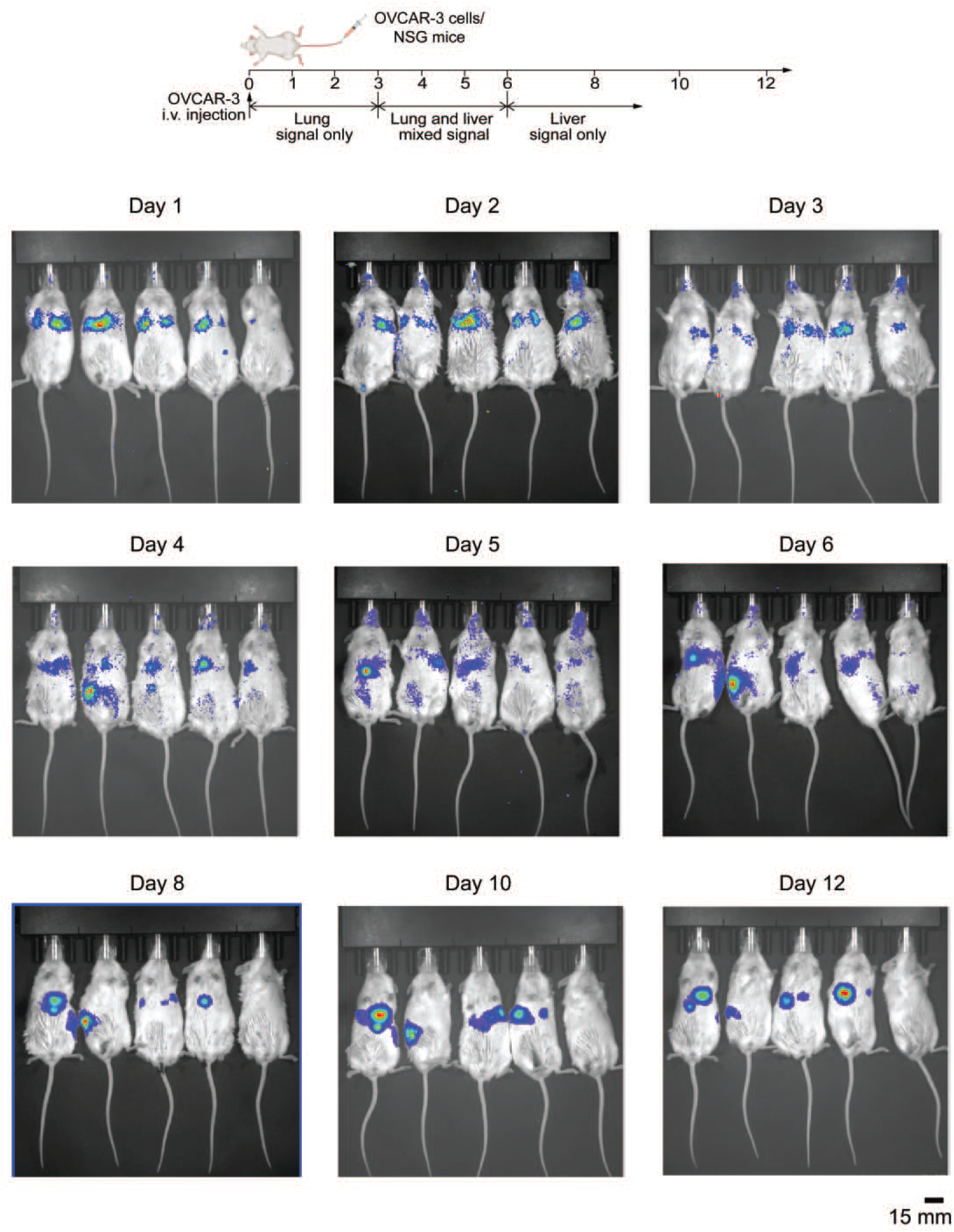
Tail vein injection of OVCAR3 ovarian cancer cells leads to establishment of liver metastases. Metastatic colonies were established in NSG mice via tail-vein injection of luciferase-expressing OVCAR-3 cells (day 0). IVIS luminescence imaging on day 1 confirmed the presence of OVCAR-3 cells in the lungs. Luminescence signal was observed exclusively in the lungs till day 3, following which, signal was observed to shift downwards to the liver (day 4-6). A week after OVCAR-3 injections, the luminescent signal was observed exclusively from the liver with no more signal being detected from the lungs. All data in this figure was generated with OVCAR-3 cells *in vivo* in NSG mice.

**Supplementary Figure 2.**
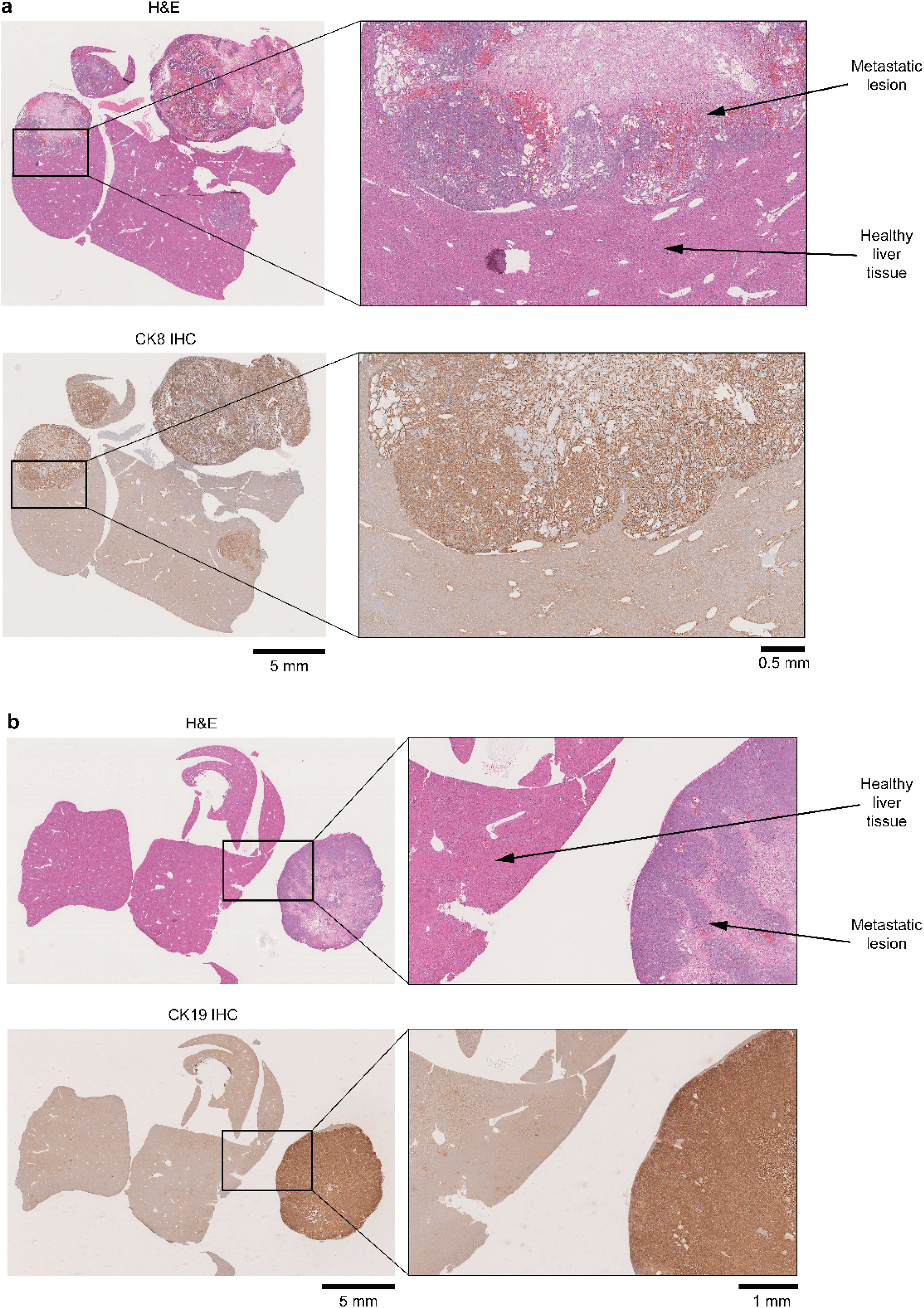
H&E-based identification of metastatic lesions is verified by cytokeratin IHC. H&E staining on liver sections showed the presence of numerous metastatic lesions on day 26. These lesions were histologically identified by their circular shape, intense purple staining, and irregular cell arrangement. Larger metastatic lesions also displayed a necrotic core indicated by lighter staining in the middle due to lack of cells. This identification of metastatic lesions was verified via IHC for **(a)** cytokeratin 8 (CK8) and **(b)** cytokeratin 19 (CK19), which are expressed abundantly by OVCAR-3 cells. All data in this figure was generated with OVCAR-3 cells *in vivo* in NSG mice.

**Supplementary Figure 3.**
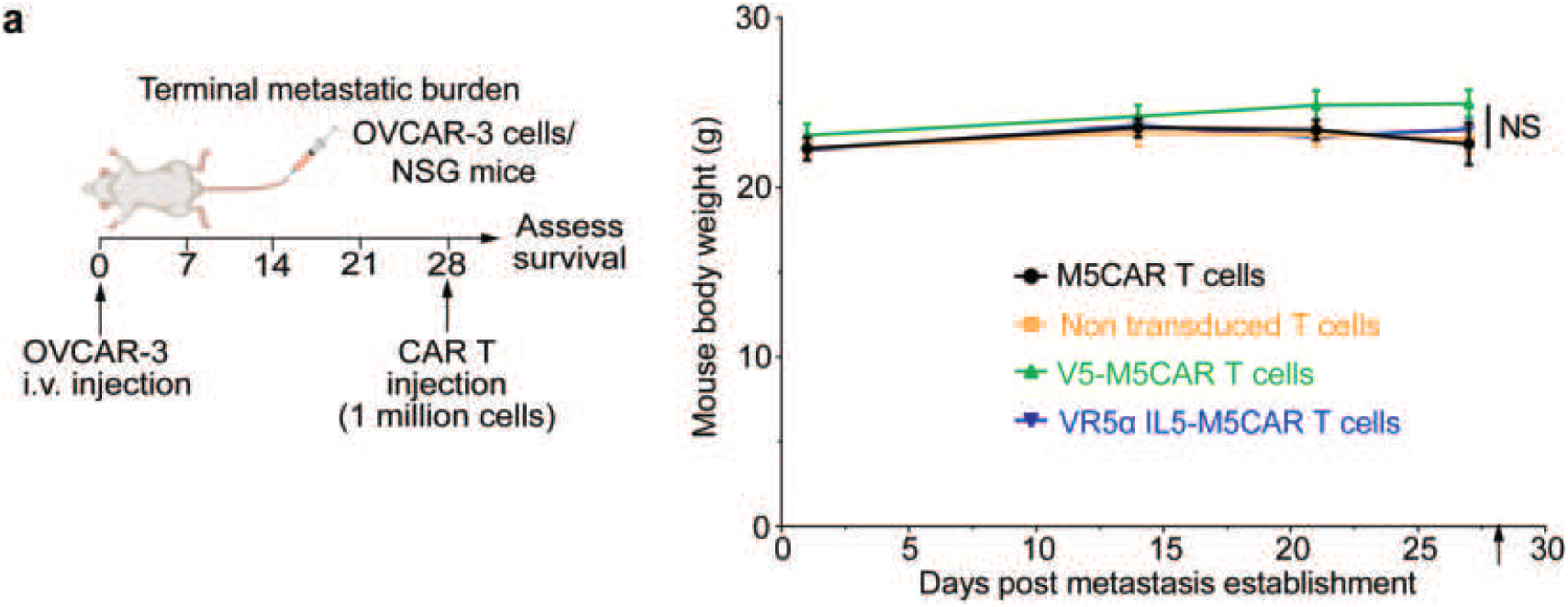
Terminal metastatic burden supplementary data. **(a)** Mouse body weights during terminal metastatic burden survival study show no significant differences during the course of the experiment. All data in this figure was generated with OVCAR-3 cells *in vivo* in NSG mice. p value: NS: Not Significant

**Supplementary Figure 4.**
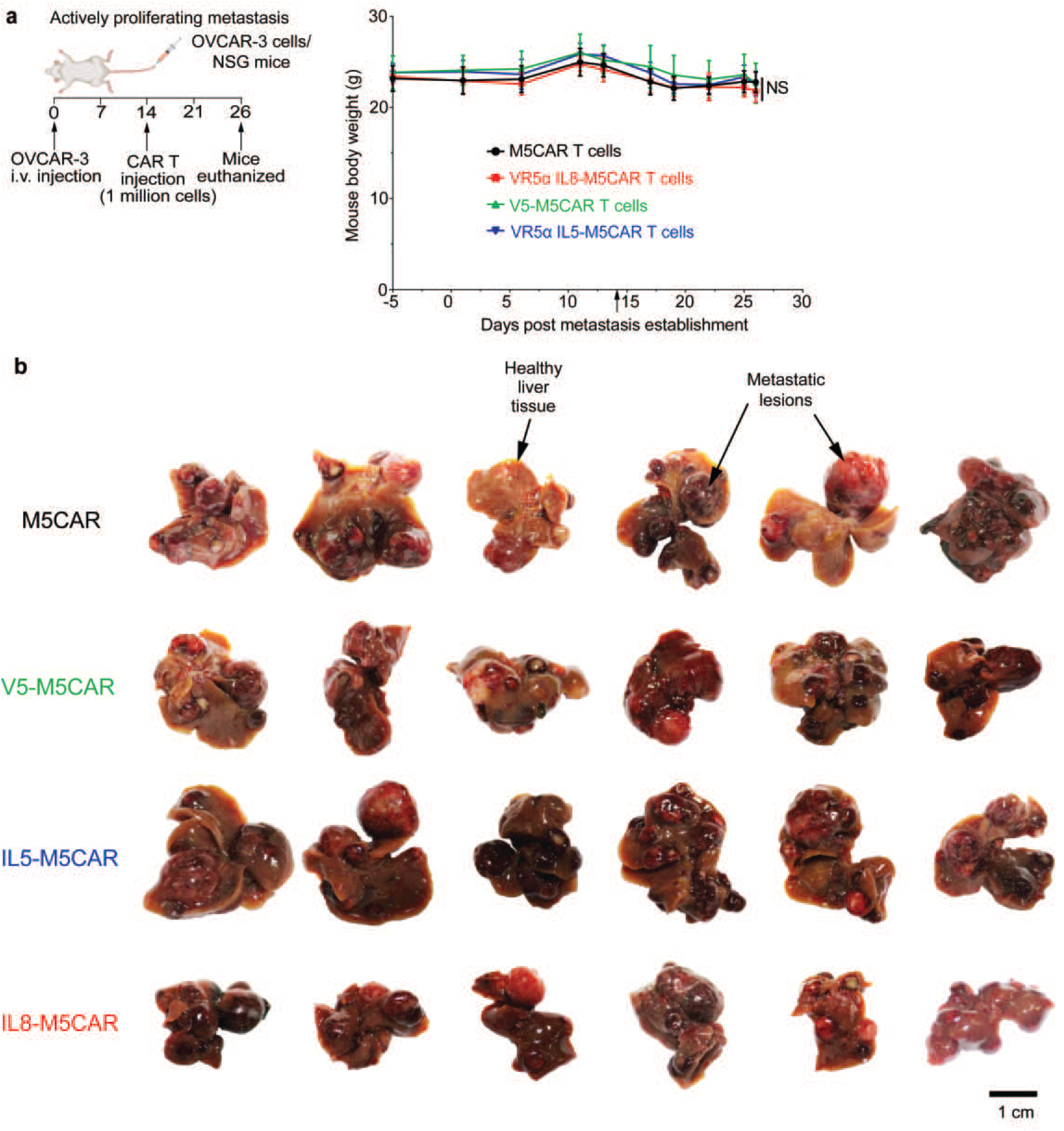
Metastatic outgrowth treatment supplementary data. **(a)** Mouse body weights during metastatic outgrowth treatment studies show no significant differences during the course of the experiment. **(b)** Images showing excised livers with metastases from all conditions. M5CAR T cell treated mice had prominent metastatic lesions, while CAR TV treated mice showed healthier livers with lesser metastasis. All data in this figure was generated with OVCAR-3 cells *in vivo* in NSG mice. p value: * p < 0.05, NS: Not Significant

**Supplementary Figure 5.**
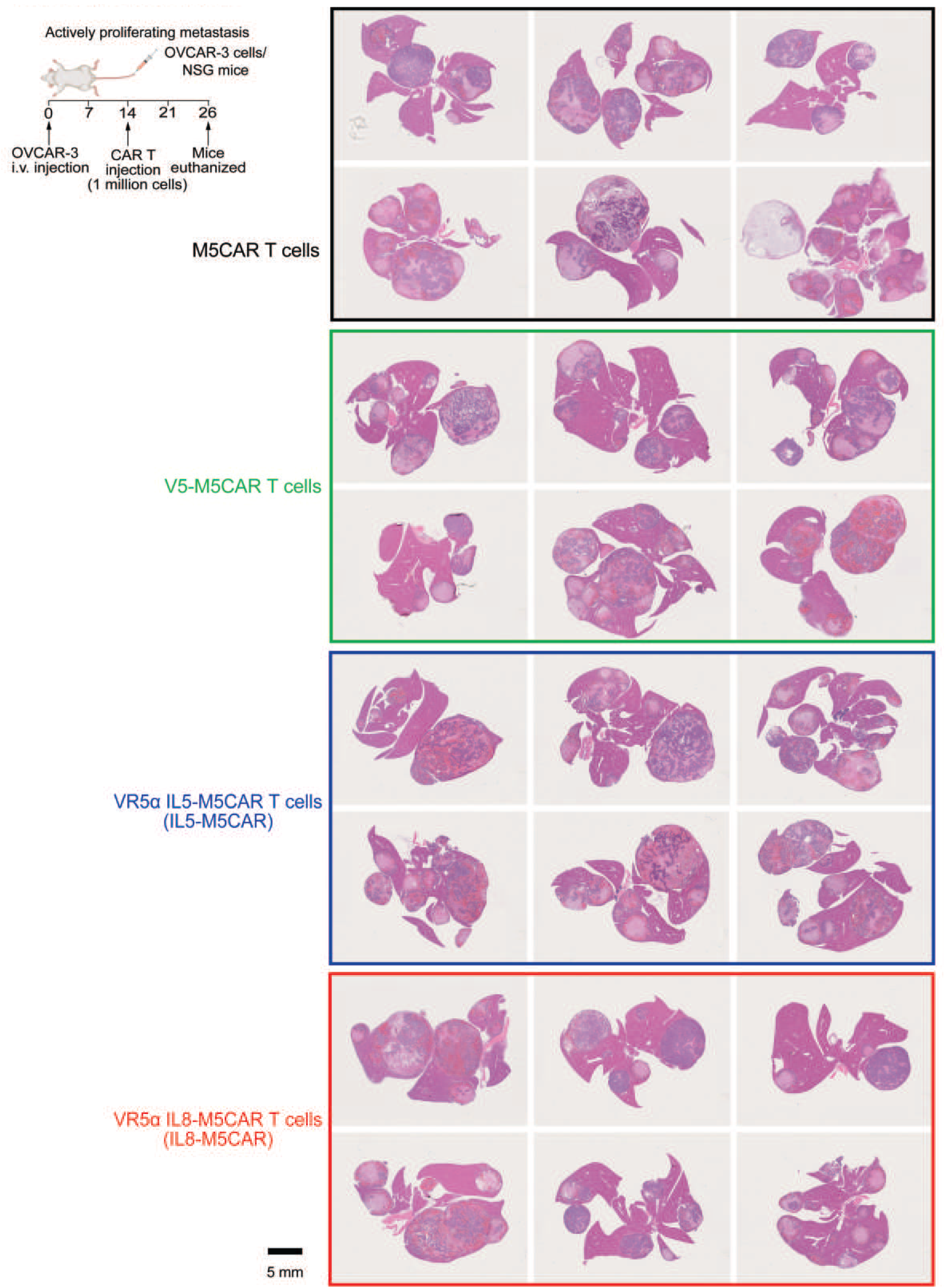
H&E sections of liver following CAR TV treatment in mice undergoing active proliferation of metastasis. Full panel of livers from mice treated with M5CAR T cells or CAR TV cells. Mice treated with CAR TV cells showed a lower metastatic burden than M5CAR T cell control, with V5-CAR being the most effective. All data in this figure was generated with OVCAR-3 cells *in vivo* in NSG mice.

**Supplementary Figure 6.**
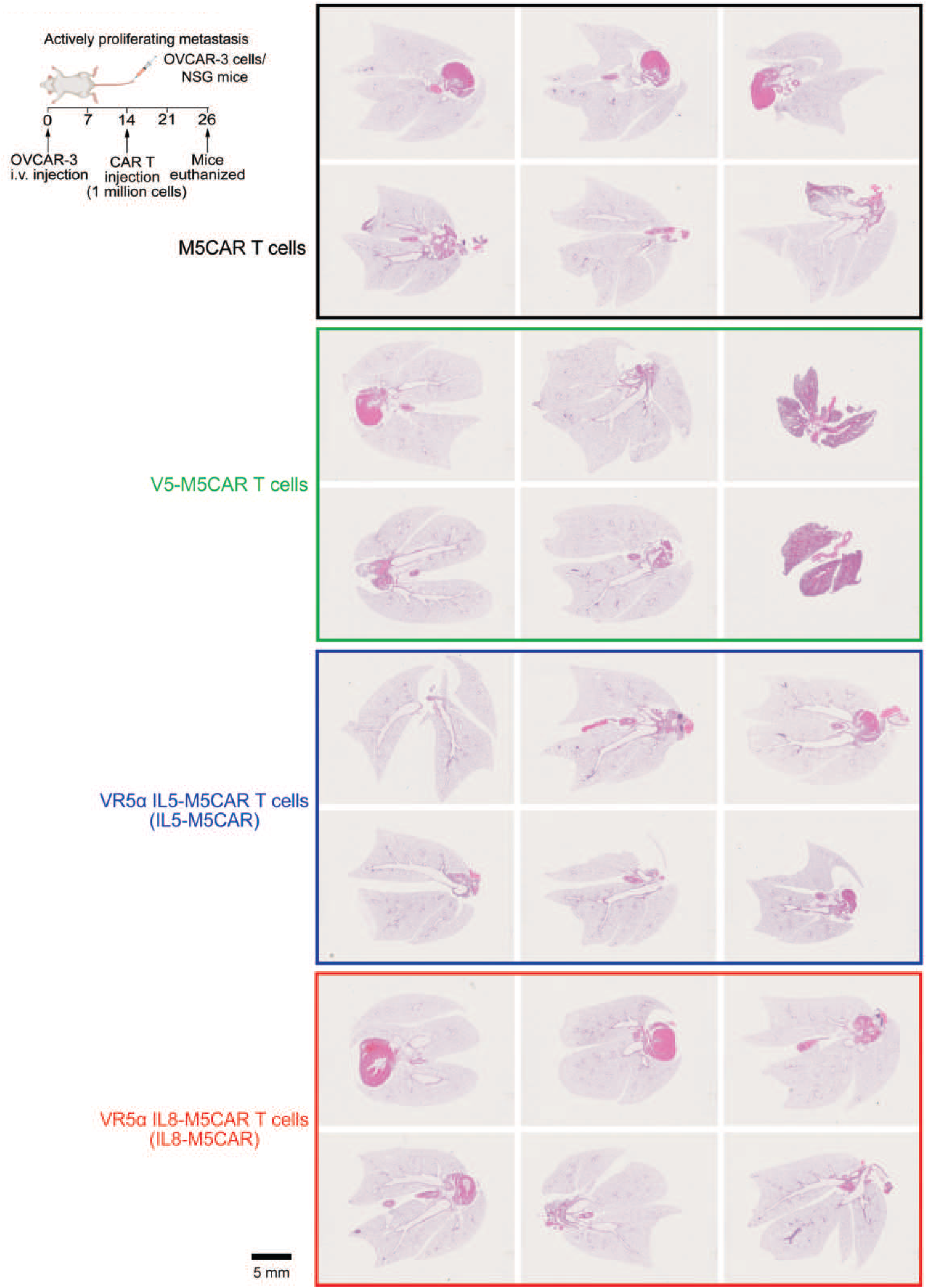
H&E sections of lung following CAR TV treatment in mice bearing actively proliferating metastasis. Full panel of lungs from mice treated with M5CAR T cells or CAR TV cells. H&E sections of lungs confirm the lack of metastatic foci, consistent with the disappearance of luminescent signal from the lungs about one week after injection. All data in this figure was generated with OVCAR-3 cells *in vivo* in NSG mice.

**Supplementary Figure 7.**
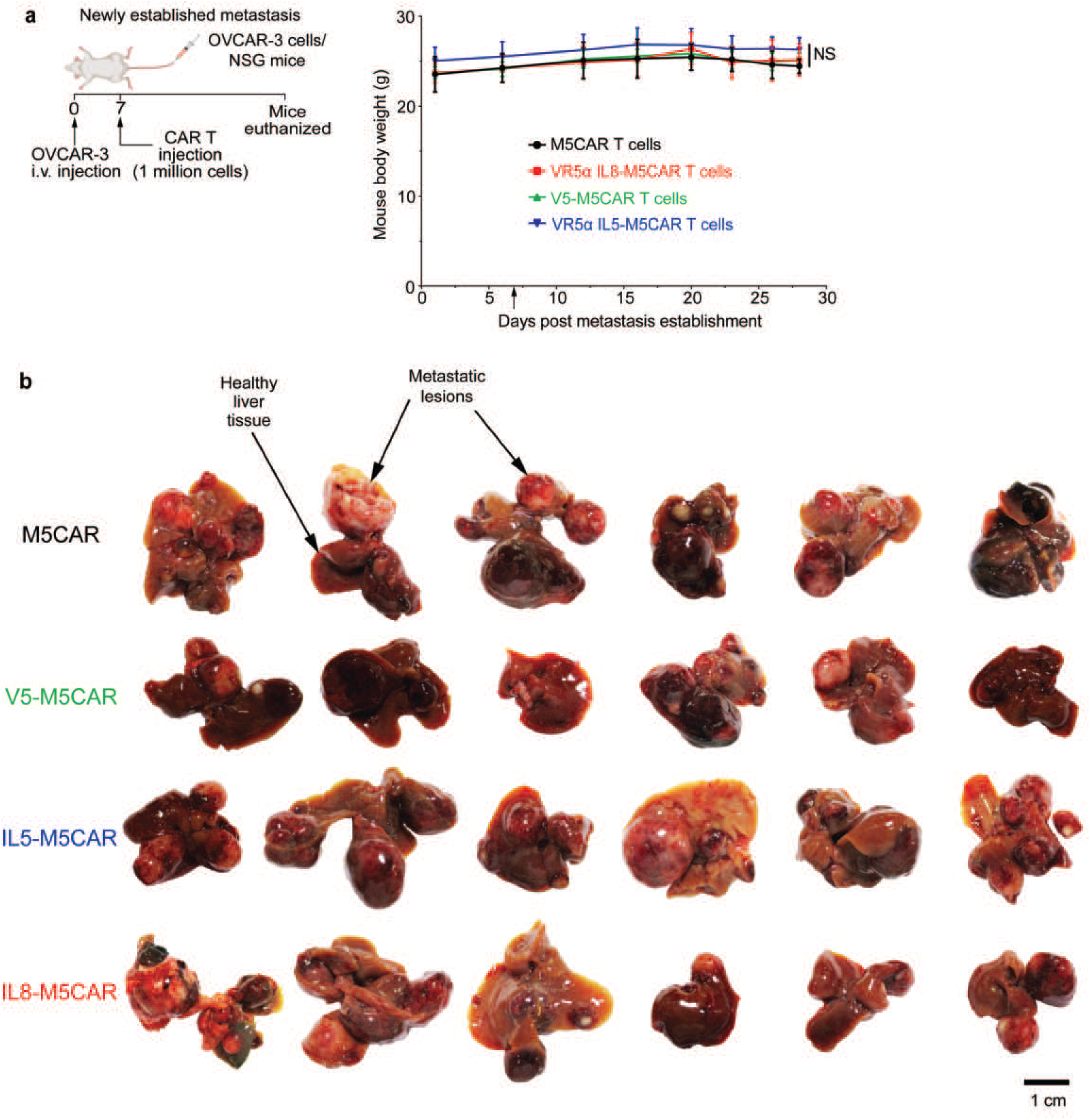
Newly-established metastasis supplementary data. **(a)** Mouse body weights show no significant differences during the course of the experiment. **(b)** Excised livers with metastases from all conditions. V5-M5CAR and IL5-M5CAR T cell treated mice showed less metastatic burden than control mice. All data in this figure was generated with OVCAR-3 cells *in vivo* in NSG mice. p value: NS: Not Significant

**Supplementary Figure 8.**
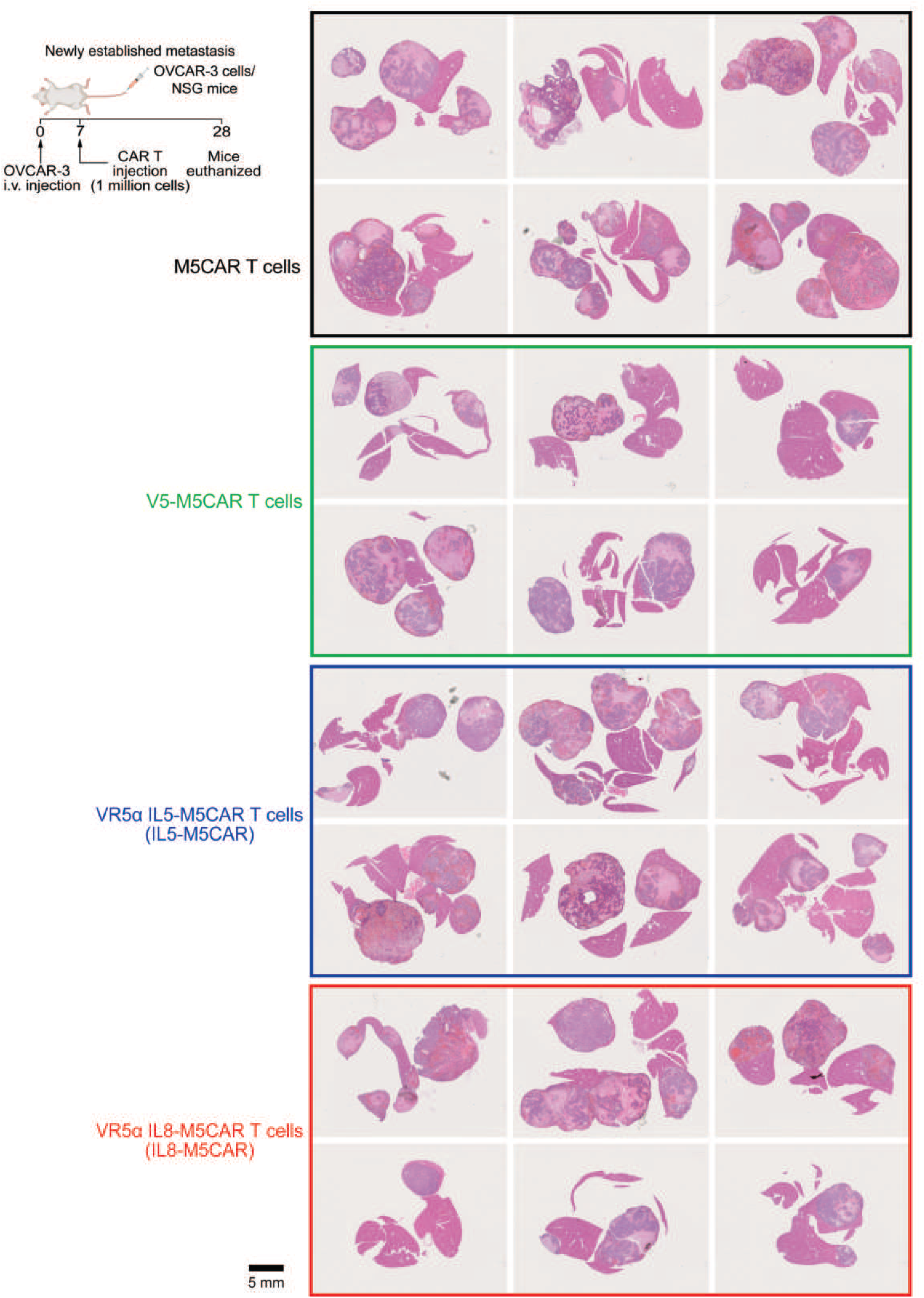
H&E sections of liver following CAR TV treatment in mice bearing newly established metastasis. Full panel of livers from mice treated with M5CAR T cells or CAR TV cells. Mice treated with CAR TV cells showed lower metastatic burden than M5CAR T cell control. All data in this figure was generated with OVCAR-3 cells *in vivo* in NSG mice.

**Supplementary Figure 9.**
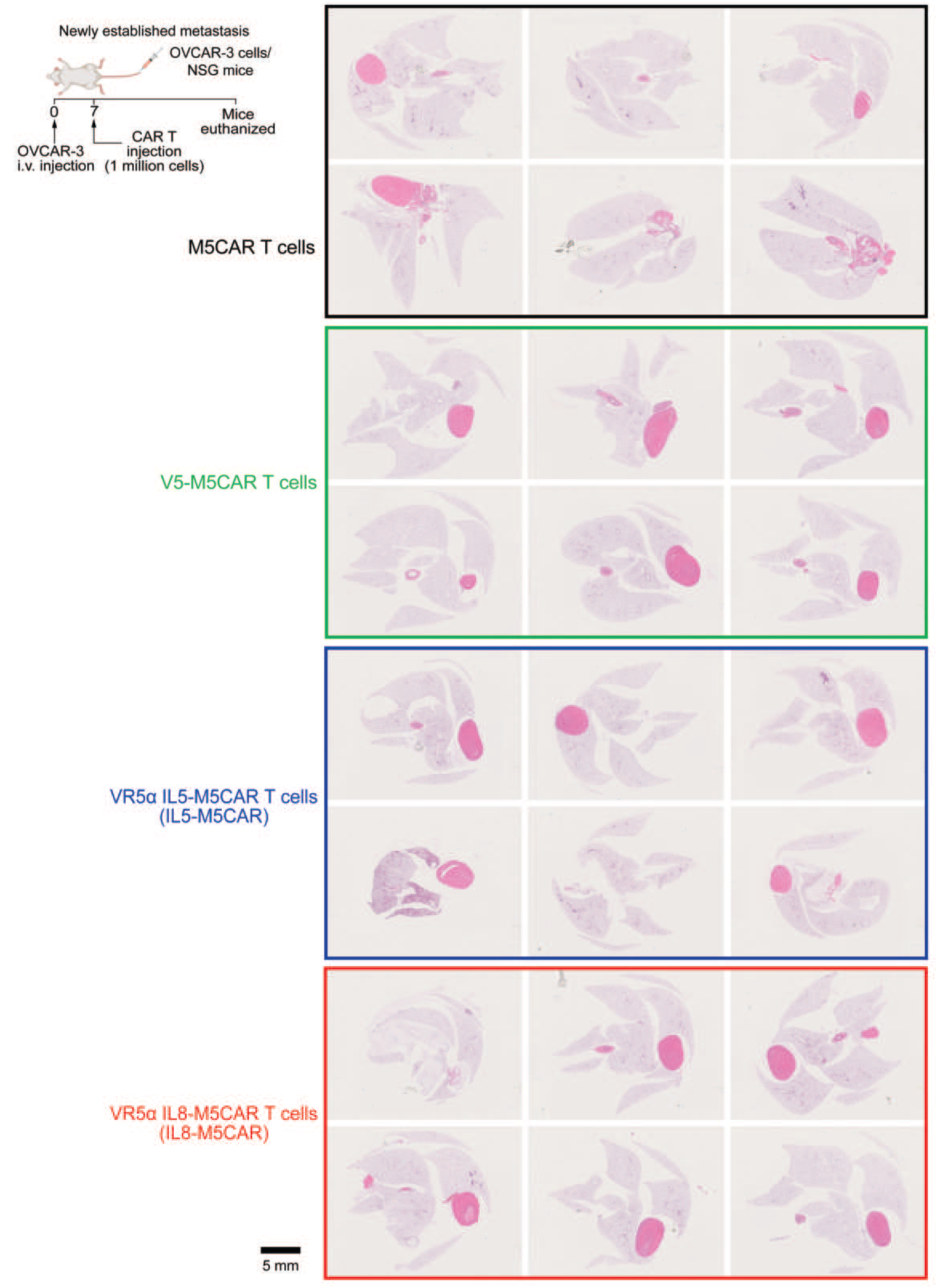
H&E sections of lung following CAR TV treatment in mice bearing newly established metastasis. Full panel of lungs from mice treated with M5CAR T cells or CAR TV cells showing the lack of any metastatic colonies. All data in this figure was generated with OVCAR-3 cells *in vivo* in NSG mice.

## Methods

### Adherent cell culture

OVCAR-3 human ovarian adenocarcinoma cells were purchased from ATCC and cultured with RPMI 1640 (Gibco) supplemented with 10% FBS, 1% penicillin/streptomycin, 200 mM L-glutamine, 1% HEPES, and 1% sodium pyruvate. These cells were cultured at 37°C and 5% CO_2_ for a maximum of ten passage numbers and tested for mycoplasma contamination every 6 months. These cells were lentivirally transduced to stably express luciferase-P2A-mCherry as described in (*50*). 293T cells were obtained from ATCC and maintained in DMEM (Corning) supplemented with 10% FBS or Opti-MEM (Gibco) supplemented with 5% FBS.

### Human T cell isolation

Primary human T cells were isolated from identity-unlinked leukapheresis packs from healthy donors who underwent same-day apheresis (Anne Arundel Medical Center, Annapolis, MD). PBMCs were isolated from leukapheresis packs immediately upon delivery using Ficoll density gradients. T cells were isolated from PBMCs using STEMCELL EasySep™ Human T Cell Isolation Kit per manufacturer’s instructions. Isolated T cells were cultured in X-Vivo 15 media (Lonza) supplemented with 5% Human AB serum, 200 mM L-glutamine, and 1% penicillin/streptomycin.

### CAR T and VR transduction

VR-GFP and M5CAR-GFP vectors were designed and produced as stated in (*50*). Lentiviral particles were generated using 293T cells. Transfection was performed with GeneJuice (Millipore Sigma), and the lentiviral particles were concentrated using Lenti-X Concentrator (Lonza). CD4 T cells (isolated using STEMCELL EasySep™ Human CD4+ T Cell Isolation Kit) and CD8 T cells (isolated using STEMCELL EasySep™ Human CD8+ T Cell Isolation Kit) were mixed in a 1:1 ratio. These were activated overnight using CD3/CD28 Dynabeads (Invitrogen) in complete X-Vivo media supplemented with 100 IU/ml IL2 (Peprotech). Activated cells were transduced with VR-GFP lentivirus overnight on Retronectin-coated plates and their transduction efficiency was verified using flow cytometry. These cells were then again transduced overnight on Retronectin-coated plates with M5CAR-GFP lentivirus, and the transduction efficiency was checked using flow cytometry. VR-expressing M5CAR, M5CAR, or non-transduced T cells were expanded for a total of 11 days. The Dynabeads were removed using Dynamags (Invitrogen) and the cells were either cryopreserved or maintained in culture in complete X-Vivo media supplemented with 100 IU/ml IL2.

### In vivo mouse modeling

All *in vivo* studies were performed in accordance with protocols approved by the Johns Hopkins University Animal Care and Use committee (ACUC). 8-12 week-old female nonobese diabetic–severe combined immunodeficient gamma (NSG) mice were purchased from Johns Hopkins Medical Institution and housed under a 12-hour dark/light cycle at 25°C. NSG mice were utilized as they are the standard model to evaluate new CAR T therapies (*73, 74*).

#### Tail vein injection to establish metastatic lesions

OVCAR-3 cells were injected via the tail vein (200,000 cells/injection) in NSG mice as per (*55, 75, 76*). 100 µl cell suspension in PBS (Gibco) was injected into the tail-vein of a mouse using a 30-gauge syringe. Mice were imaged the next day to confirm the presence of cancer cells (see below).

#### IVIS bioluminescence imaging

Mice were injected intraperitoneally (*77*) with 150 µl of 20 mg/ml D-luciferin (Goldbio, Sodium salt) solution in PBS immediately prior to anesthesia. IVIS imaging was performed per (*78*). Mice were anesthetized and maintained via inhalation of 1.5% isoflurane in 1.5 L/min oxygen. Images were acquired using a PerkinElmer IVIS Spectrum optimized for luminescence imaging. The following parameters were used for IVIS luminescent imaging: Imaging.Mode value - P+L, Photo.Exposure value – Auto, Photo.Binning value – Medium, Photo.FStop value – 8, Exposure value – 0.3 to 120 s, Binning value – Medium, FStop value – 1, FOV value – D, Height value -1.50. For the images to be suitable for quantification, the exposure time was varied according to signal to avoid images with saturated pixels. All mice were imaged between 15-30 minutes after the luciferin injection in order to keep the luminescent signal steady. Luminescent, background, and photograph images were captured, and the overlays were analyzed and presented in figures. Mice were imaged every 3-4 days from the day after metastasis establishment till the end of the experiment. Analysis of luminescent intensity was done using Living Image (PerkinElmer). Total photon flux (p/s) was analyzed, and the final values were calculated by subtracting the mean background value from the raw signal value.

#### CAR TV treatment

At a predetermined timepoint (28 days for terminal metastasis burden, 14 days for proliferating metastasis burden, or 7 days for newly established metastasis burden), mice were administered CAR TV cells via a tail-vein injection of 1 million cells in 100 µl PBS. M5CAR T cells or non-transduced T (NTD) cells that did not express VRs were the negative control. For mice being treated at a terminal metastatic burden, bioluminescence imaging was conducted one day prior to the treatment to confirm the presence of an extremely high metastatic burden. Mouse body weight was monitored every 3-4 days. Mice were sacrificed at predetermined time points as indicated for IVIS-based monitoring of metastasis. For survival experiments, mice were imaged and monitored regularly. When the luminescent signal was calculated to be above a set threshold defined in the ACUC protocol or if the mouse showed any signs of distress, the mouse was euthanized, and an event was marked. The liver and agarose perfused lungs were excised, fixed, paraffin embedded, H&E stained, and imaged as described in (*79*).

#### Immunohistochemistry

Immunohistochemistry (IHC) was performed at the Oncology Tissue and Imaging Services Core of Johns Hopkins University School of Medicine.

##### CK8

Immunolabeling for CK8 was performed on formalin-fixed, paraffin embedded sections on a Ventana Discovery Ultra autostainer (Roche Diagnostics). Briefly, following dewaxing and rehydration on board, epitope retrieval was performed using Ventana Ultra CC1 buffer (catalog# 6414575001, Roche Diagnostics) at 96 °C for 64 minutes. Primary antibody, anti-CK8 (1:2000 dilution; catalog# ab53280, Abcam) was applied at 37 °C for 60 minutes. Primary antibodies were detected using an anti-rabbit HQ detection system (catalog# 7017936001 and 7017812001, Roche Diagnostics) followed by Chromomap DAB IHC detection kit (catalog # 5266645001, Roche Diagnostics), counterstaining with Mayer’s hematoxylin, dehydration and mounting.

##### CK19

Immunolabeling for CK19 was performed on formalin-fixed, paraffin embedded sections on a Ventana Discovery Ultra autostainer (Roche Diagnostics). Briefly, following dewaxing and rehydration on board, epitope retrieval was performed using Ventana Ultra CC1 buffer (catalog# 6414575001, Roche Diagnostics) at 96 °C for 64 minutes. Primary antibody, anti-CK19 (1:400 dilution; catalog# ab7754, Abcam) was applied at 37 °C for 60 minutes. Primary antibodies were detected using an anti-mouse HQ detection system (catalog# 7017936001 and 7017782001, Roche Diagnostics) followed by Chromomap DAB IHC detection kit (catalog # 5266645001, Roche Diagnostics), counterstaining with Mayer’s hematoxylin, dehydration and mounting.

### CellViT++ analysis

Python (version 3.10) was used to perform cell segmentation and classification on digitized H&E slides. Cell segmentation was carried out using CellViT++, a transformer-based model trained on liver samples to ensure optimal performance on our dataset (*66, 67*). Cell classification was performed using the SAM-H classifier, reported by Horst et al. (*66, 67*) as the best-performing model for histopathology cell type prediction. Each cell was classified into one of five categories: connective, dead, epithelial, inflammatory, or neoplastic. CellViT++ produced GeoJSON outputs containing cell contours and cell type annotations, from which bulk counts by cell type were derived.

To analyze the metastasis regions, the H&E slides were processed at OpenSlide level 2 (corresponding to approximately 5× resolution for 20× whole-slide images) to generate metastasis masks. Metastatic regions were identified using HSV color thresholding (H: 110–160) combined with value-channel filtering (V < 215) to isolate dark hematoxylin-rich regions. Morphological opening and closing operations with a 5×5 kernel was applied to remove noise and fill small gaps, resulting in a clean binary tumor mask. Inflammatory cells (type = 2) from the CellViT++ output were then spatially filtered to retain only those whose centroids fell within the tumor mask. This provided a count of inflammatory cells specifically within metastatic regions.

For visualization, tumor masks and inflammatory cell centroids were overlaid on the corresponding slide thumbnails and saved as image files. Counts per slide were tabulated and exported to CSV for downstream analysis.

### Statistics

All *in vivo* experimental conditions were performed with a minimum of 5 mice bearing one tumor each (N=5). *A priori* power analysis was performed using t-tests in G*Power 3.1.9.2 software using the following parameters: alpha as 0.05, power of 0.65, allocation ratio (N2/N1) as 1, effect size (ρ) as large, and analysis with one-tailed t-test. Mice were randomized prior to initiation of treatment. For survival study, significance, hazard ratios, and 95% confidence intervals were calculated via logrank (Mantel–Cox) test. Other experiments were analyzed using one-way ANOVA (wherever number of conditions ≥ 3). After confirming significance by ANOVA, significance between select key conditions were also analyzed by two-tailed t-test. p-values <0.05 was considered significant (*p<0.05, **p<0.01, ***p<0.001, ****p<0.0001, NS: Not Significant). In this manuscript, the usage of the word “significant” and its variations refer to statistical significance. All figure plotting and statistical calculations were performed using GraphPad Prism.

## Materials availability

This study did not generate new unique reagents.

## Data availability

No large scale data was generated in this manuscript.

## Author’s Contributions

P.R.N. and D.W. conceived the project. P.R.N., E.A.H., C.B., and D.W. designed the experiments. P.R.N. and E.A.H. performed the experiments. P.R.N. and S.J. analyzed the data. P.R.N. wrote the manuscript, which was read and edited by all authors.

## Acknowledgements

This work was supported by grants from the National Cancer Institute (U54CA143868, U54AR081774, and U54CA268083) and the National Institute on Aging (U01AG060903) to D.W; and P30 CA006973 (to Oncology Tissue and Imaging Services Core of Johns Hopkins University School of Medicine)

